# An actin nucleation complex catalyzes filament formation at sites of exocytosis

**DOI:** 10.1101/715409

**Authors:** Oliver Glomb, Yehui Wu, Lucia Rieger, Diana Rüthnick, Medhanie Mulaw, Nils Johnsson

**Affiliations:** Institute of Molecular Genetics and Cell Biology, Department of Biology, Ulm University, James-Franck-Ring N27, D-89081 Ulm, Germany

**Keywords:** polar growth, vesicular traffic, actin nucleation, cdc42

## Abstract

Due to the local enrichment of factors that influence its formation, dynamics, and organization, the actin cytoskeleton displays different shapes and functions within the same cell. In yeast cells post-Golgi vesicles ride on long actin cables to the bud tip. The scaffold proteins Boi1 and Boi2 participate in tethering and docking these vesicles to the plasma membrane. Here we show that Boi1/2 also recruit nucleation and elongation factors to form actin filaments at sites of exocytosis. Disrupting the physical connection between Boi1/2 and the nucleation factor Bud6 impairs filament formation in the bud, reduces the directed movement of the vesicles to the tip, and shortens their tethering time at the cortex. Artificially transplanting Boi1 from the bud tip to the peroxisomal membrane partially redirects the actin cytoskeleton and the vesicular flow towards the peroxisome, and creates an alternative, rudimentary vesicle-docking zone. We conclude that Boi1/2 is sufficient to induce the formation of a cortical actin structure that receives and aligns incoming vesicles before fusing with the membrane.

## Introduction

The directed transport of post-Golgi vesicles influences the shape of cells and forms diverse structures such as axons in animals, hyphal extensions in fungi, or pollen tubes in plants. Polar growth is the result of a complex interplay between RhoGTPase-based signaling, cytoskeletal organization, vesicular traffic and fusion. By growing through budding, yeast serves as a model organism to understand the molecular mechanisms behind polar growth. The bud tip of yeast cells is the site of preferred exocytosis and hosts a complex assembly of proteins that tether the vesicle to the plasma membrane. Vesicles arrive at these sites on actin cables by a myosin type-V (Myo2) driven transport (Jin et al. 2011). Post-Golgi vesicles do not immediately fuse with the PM but stay immobile for a defined time at or close by the site of their prospective fusion. This dwell time depends among other factors on the vesicle-bound Myo2 and thus on the presence of an actin structure adjacent to the site of fusion (Donovan and Bretscher 2015; Lepore, Martinez-Nunez, Munson 2018).

Linear actin cables that point to the bud tip are formed by the nucleation-promoting factor Bud6 in cooperation with the formin Bni1 (Amberg et al. 1997; Graziano et al. 2011; Moseley et al. 2004). Bud6 and Bni1 co-localize to the tip of the bud as members of the polarisome multi-protein complex (Amberg et al. 1997; Sheu et al. 1998). Full activation of this complex requires the binding of small Rho-GTPases to relive the auto-inhibition of Bni1 (Evangelista et al. 1997; Li, F. and Higgs 2003; Li, F. and Higgs 2005). Deletion of core components of the polarisome dissolve the focused distribution of Bud6 and Bni1, yet leave actin filament formation and the delivery of vesicles less tip-directed but otherwise still largely intact (Tcheperegine, Gao, Bi 2005). This finding suggests the existence of additional factors that position filament formation at the membrane of the bud. We describe a complex consisting of Bud6, Bni1 and the vesicle tethering factors Boi1 and Boi2 that initiates the formation of actin filaments at or close to the sites of vesicle fusion.

## Results

### Boi1/2 physically interact with actin nucleation and elongation factors

To identify alternative regulators of actin filament nucleation in the bud, we performed a systematic Split-Ubiquitin (Split-Ub) interaction analysis and screened Bni1 and Bud6 as C_ub_-RUra3-fusions (CRU) against an array of 504 N_ub_ fusion proteins (Hruby et al. 2011; Johnsson and Varshavsky 1994; Wittke et al. 1999). The Split-Ub analysis revealed among specific binding partners for Bni1 or Bud6, Boi1 and Boi2 as shared interaction partners of both proteins (Fig. 1A, Fig. S1, Table S1). Boi1 and Boi2 are homologous proteins that assist in the tethering and fusion of post-Golgi vesicles at the plasma membrane and thus qualify as candidates for organizing actin filaments in the bud (Kustermann et al. 2017; Masgrau et al. 2017). Both proteins were already shown to interact with Bud6 *in vivo* (Kustermann et al. 2017). As Boi1 and Boi2 perform their essential functions redundantly, we performed our molecular analysis predominantly on Boi1 (Kustermann et al. 2017). Split-Ub interaction analysis in strains lacking either *BUD6* or *BNI1* confirmed that both proteins interact independently of each other with Boi1 (Fig. 1B). N_ub_ fusions to fragments of Boi1 or Boi2 localized the binding sites for Bud6 and Bni1 to their N-terminal 300 residues (Fig. 1C.) Split-Ub analysis of CRU fusions to fragments of Bni1 placed the binding site for Boi1 within the N-terminal 854 residues of Bni1 and thus away from the C-terminally located binding site for Bud6 (Fig. 1D). N_ub_ fusions to fragments of Bud6 allowed confining the Boi1-interaction site to the N-terminal 141 residues of Bud6 (Fig. 1E). We expressed the so-defined minimal binding fragments of Boi1 (1-203), Bud6 (1-141), and Bni1 (1-854) in *E.coli* and could show by pull down analysis that the interaction between Boi1 and Bni1 and between Boi1 and Bud6 are direct (Fig. 1F). To obtain a mutation that disrupts the interaction to Bud6 without grossly disturbing the structure of Boi1, we further fine-mapped the Bud6 interaction site on Boi1. A screen of N_ub_ fusions to different N-terminal fragments of Boi1 identified the linker region (residues 77-178) between the SH3 and the SAM domain as autonomous binding site for Bud6 (Fig. S2). A pull-down of the purified His-tagged Bud6_1-141_ with a GST fusion to Boi_77-178_ confirmed our analysis (Fig. S2). We deleted the Boi1-linker region in the yeast genome (*boi1*_Δ86-178_) and tested the Boi1_Δ86-178_CRU-expressing strain against the N_ub_ array. The analysis confirmed that Boi1_Δ86-178_ had specifically lost its interaction to Bud6 (Fig. 1G). The interactions with other polarity proteins like Bem1 as well as Sec1 remained unaffected by this deletion (Figs. 1G, S2). The deletion did not detectably impair the interaction between Boi1 and Bni1 (Fig.1H). Overexpressing Boi1-mCherry leads to large un-budded cells whose plasma membrane is stained by Boi1-mCherry (Fig. 1J). Co-staining the actin structures in these cells with Lifeact, or co-staining with GFP fusions to Bud6 or Bni1 visualizes the relocation of all three proteins to the Boi1-mCherry-stained cortex, suggesting that Boi1 binds to the Bni1/Bud6 complex in its active actin filament-promoting conformation (Figs. 1I, J).

**Figure1.**
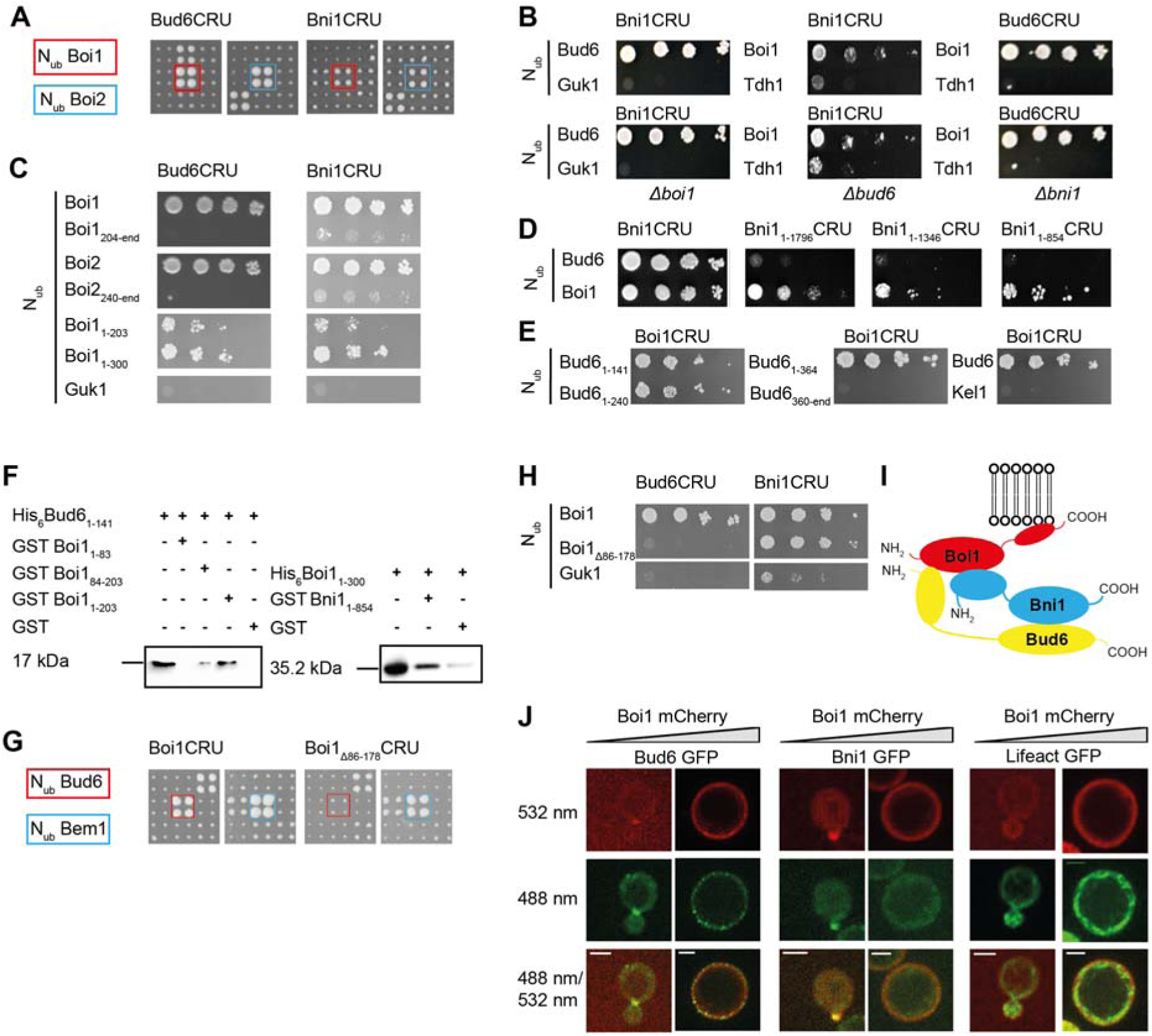
Boi1/2 interact with Bud6 and Bni1. (A) Cut-outs of a Split-Ub array of diploid yeast cells containing Bud6CRU (left panels), or Bni1CRU (right panels), and co-expressing different N_ub_ fusions. The N_ub_- and CRU-expressing cells were independently mated four times, spotted in quadruplets and transferred onto medium containing 5-FOA. Growth of four colonies indicates interaction. N_ub_-Boi1, and N_ub_-Boi2 are highlighted in red and blue (See Fig. S1 for complete array, and Table S1 for list of interaction partners). (B) 4 µl of yeast cultures co-expressing the indicated N_ub_- and C_ub_ fusion proteins were spotted in 10-fold serial dilutions, starting with an OD_600_ = 1, on medium containing 5-FOA. N_ub_ fusions to Guk1, or Tdh1 should not interact. Upper lanes show the interactions in otherwise wild type yeast. Lower lanes show interactions in the absences of *BOI1* (left panel), *BUD6* (middle panel) or *BNI1* (right panel). (C) As in (B) but with cells co-expressing Bud6CRU (left lanes) or Bni1CRU (right lanes) and the indicated fragments of Boi1 or Boi2 as N_ub_ fusions. (D) As in (B) but with cells co-expressing N_ub_-Boi1 or N_ub_-Bud6 and C-terminally truncated fragments of Bni1 as CRU fusions. (E) As in (B) but with cells co-expressing Boi1CRU together with Bud6 or fragments of Bud6 as N_ub_ fusions. N_ub_-Kel1 should not interact. (F) Extracts of *E. coli* cells expressing 6xHis fusion of Bud6_1-141_ (left panel), or Boi1_1-300_ (right panel) were incubated with sepharose beads exposing GST, or GST-fusions to fragments of Boi1 (left panel), or Bni1 (right panel). Input and bound fractions were separated by SDS-PAGE and analyzed by anti-His antibodies after transfer onto nitrocellulose. Corresponding SDS-PAGE and quantification of the Western-blots are provided in Figure S2A-D. (G) as in (A) but with cells expressing Boi1CRU or Boi1_Δ86-178_CRU. Interactions of N_ub_-Bud6 and N_ub_-Bem1 are highlighted in red and blue (See Fig S2 for complete arrays). Bud6 does not interact with Boi1_Δ86-178_. (H) as in (B) but with cells co-expressing Bud6CRU (left panel), or Bni1CRU (right panel) with the indicated N_ub_ fusions. Boi1_Δ86-178_ still interacts with Bni1. (I) Cartoon summarizing the *in vivo* and *in vitro* interaction data. The interaction between Bud6 and Bni1 was extensively studied by others (Graziano et al. 2011; Tu et al. 2012). See results section for details. (J) Yeast cells containing *P_GAL1_-BOI1-mCherry* and co-expressing either Bud6-GFP, Bni1-GFP, or Lifeact were shifted to galactose for 4h (left panels) and 24h (right panels) to induce overexpression of Boi1-mCherry. Shown are the mCherry channel (upper row), the GFP channel (middle row) and the overlay of both channels (lower row). All the GFP fusions are recruit to the cortex upon Boi1 overexpression. Scale bars indicate 2 µm.

### The Boi proteins influence the actin cytoskeleton independently of their vesicle fusion activity

Boi1/2 might guide the Bud6/Bni1-complex to generate actin filaments at sites of exocytosis. We visualized the actin cytoskeleton in Δ*boi1*Δ*boi2* cells expressing fragments of Boi1 of increasing length. The C-terminal PH domain of Boi1 is the minimal fragment that rescues the essential function of Boi1/2 during vesicle fusion. Cells expressing this domain (*boi1*_Δ_*_414_* Δ*boi2)* were significantly enriched in delocalized actin patches and contained fewer and thinner actin cables as the corresponding *BOI1*Δ*boi2-*cells (Fig. 2A, B). This effect documents a clear impact of Boi1/2 on the actin cytoskeleton. We next correlated the presence of the Bud6/Bni1 binding sites on the expressed Boi1-fragments with changes of the actin cytoskeleton in these cells. A deletion of the first 203 residues removes the major Bud6 binding site of Boi1 and affects the binding to Bni1. This deletion already reduced the percentage of actin cables and also increased the amount of de-localized actin patches. Additionally removing the SAM domain of Boi1 (*boi1*_Δ_*_299_*) and thus any residual interactions with the Bud6-Bni1 complex further lowered the amount of actin cables and enhanced the number of delocalized actin patches to similar levels as found in cells expressing only *Boi1*_Δ_*_414_*. The impaired vesicle fusion of a Δ*boi1*Δ*boi2* strain can be suppressed by overexpression of the t-SNARE Sso1 (Kustermann et al. 2017). The still disorganized actin structure of this strain confirms that actin organization and vesicle docking are distinct activities of Boi1/2 (Fig. 2B). A GFP fusion to the Rab-GTPase Sec4 is a marker for post-Golgi vesicles, which become highly polarized in the growing bud (Jin et al. 2011). Truncating Boi1 from its N-terminus increasingly dissolved the tip-focused distribution of these vesicles (Fig. 2A, C). This observation further substantiates the role of Boi1/2 in actin organization as secretory vesicles strictly travel on actin cables (Pruyne, Schott, Bretscher 1998).

**Figure 2.**
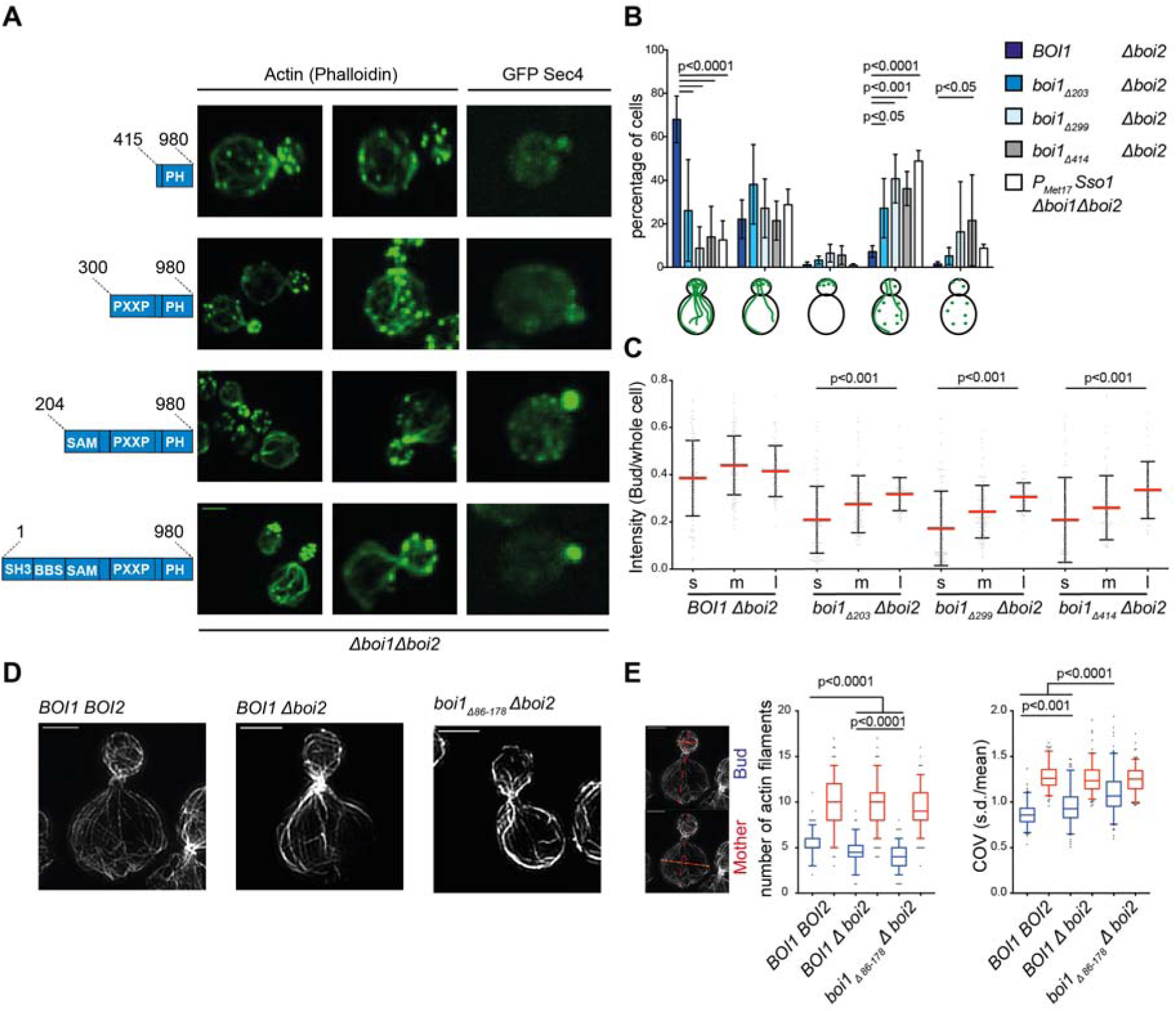
Mutations in Boi1 influence the actin cytoskeleton. (A) Fluorescence microscopy of Δ*boi2*-cells expressing different, genomically integrated fragments of *BOI1* under a copper inducible promoter after staining actin with Phalloidin-Alexa488 (left and middle panel), or co-expressing GFP-Sec4 as marker for post-Golgi vesicles (right panel). Scale bar indicates 2 µm. (B) Quantification of actin phenotypes of cells in (A) according to the categories as shown by the cartoons in the lower panel. Analysis was based on three independent experiments with in total n*_BOI1,_* _Δ_*_boi2_=642,* n*_boi1_*_Δ_*_203,_* _Δ_*_boi2_*=*823,* n*_boi1_*_Δ_*_299,_* _Δ_*_boi2_*=*333,* n*_boi1_*_Δ_*_414,_* _Δ_*_boi2_*=666 and n *_PMET17-SSo1_*_Δ_*_Boi1_*_Δ_*_414,_* _Δ_*_boi2_*=257 cells. Statistical analysis was performed with a two-way ANOVA using a Tukeys-Post-test for multiple comparisons. (C) The ratio of the GFP-Sec4 fluorescence intensities between mother and bud were used to quantify the degree of polarized secretory vesicles in small (s, <1.5 µm), medium (m, 1.5-3 µm) and large buds (l, > 3 µm) of the cells in (A). The experiments were performed in duplicates with two clones analyzed for each genotype resulting in a total cell number of n*_BOI1,_* _Δ_*_boi2_=*114(s), 135(m), 63(l), n*_boi1_*_Δ_*_203,_* _Δ_*_boi2_*=107(s), 118(m), 65(l), n*_boi1_*_Δ_*_299,_* _Δ_*_boi2_*=101(s), 133(m), 52(l), n*_boi1_*_Δ_*_414,_* _Δ_*_boi2_*=68(s), 103(m), 51(l). Statistical analysis with calculated p-values are based on a non-parametric Kruskal-Wallis test followed by Dunńs post test for multiple comparisons. (D) Representative images of cells, treated with 1µM CK-666 for 10 min to dissolve actin patches, were subsequently fixed, stained with Alexa488-Phalloidin, and inspected with SIM. Scale bar indicates 2 µm. (E) SIM images of in total n*_BOI1, BOI2_*= 149(tip), 175(mother), n*_BOI1,_* _Δ_*_boi2_*= 150(tip), 199(mother), n*_boi1_*_Δ_*_86-178,_* _Δ_*_boi2_*= 169(tip), 220(mother) cells were analyzed from three independent measurements to quantify the number of actin cables crossing a virtual plane in bud (blue) and mother cell (red) (left, and middle panel). In total n*_BOI1, BOI2_*=123 (tip, mother), n*_BOI1,_* _Δ_*_boi2_*= 135 (tip, mother) and n*_boi1_*_Δ_*_86-178,_* _Δ_*_boi2_*= 167 (tip, mother) cells were taken to measure the actin cable densities (right panel) (COV= coefficient of variation). Statistical analysis was performed with a Kruskal-Wallis test followed by Dunńs post-test for multiple comparisons. All error bars show the s.d of the mean.

### The Boi1-Bud6 complex stimulates actin filament formation in the bud

The high density of actin patches interferes with the simultaneous detection of actin cables in the bud. We thus repeated the actin staining of Δ*boi2*-cells carrying the minimally perturbed *boi1*_Δ_*_86-178_* allele after treatment with the actin patch inhibitor CK-666 (Hetrick et al. 2013). Structured illumination microscopy (SIM) of the actin cytoskeleton showed that the bud of *boi1*_Δ_*_86-178_* Δ*boi2* cells was less densely filled with actin filaments than *BOI1* Δ*boi2* cells (Fig. 2D). Applying the coefficient of variation as quantitative measure we could support the conclusion derived from the visual analysis of the cells (Fig. 2E) (Garabedian et al. 2018)(see Materials and Methods). Compared to wild type *BOI1BOI2-* and *BOI1*Δ*boi2*-cells, *boi1*_Δ_*_86-178_* Δ*boi2-*cells also displayed fewer actin filaments crossing a virtual plane that was placed in the bud perpendicular to the polarity axis (Fig. 2E). In contrast, actin filament number and density were not changed in mother cells upon deletion of the Bud6 binding site in Boi1 (Fig. 2E, F). The polar localization of Bud6 was not affected in *boi1*_Δ_*_86-178_* Δ*boi2-*cells (Fig. S3).

### Cortical actin modulates vesicular flow and fusion

To measure the influence of the Boi1/2 induced actin structures on the movement and fusion of post-Golgi vesicles, we photo-bleached the buds of cells expressing GFP-Sec4 and compared the trajectories of individual vesicles entering the bud of *BOI1 BOI2-*, *BOI1* Δ*boi2-* and *boi1*_Δ_*_86-178_* Δ*boi2*-cells by time-lapse microscopy (Video_S1) (Donovan and Bretscher 2015). Incoming vesicles in wild type cells were often directly transported to the cell cortex, where they either tethered and subsequently fused, or moved along the cortex to the tip where fusion occurred. Incoming vesicles in Δ*boi2-* and more pronounced in *boi1*_Δ_*_86-178_* Δ*boi2*-cells took longer to find their final destination (Video_S1). Especially post-Golgi vesicles in *boi1*_Δ_*_86-178_* Δ*boi2*-cells headed after their first contact with the cortex to a different region of the cortex or tumbled in the center of the bud. We occasionally observed an incoming vesicle that was redirected into the mother cell after shortly touching the cortex of the bud. Moreover, incoming vesicles seemed to reside longer at the neck before entering the bud (Video_S1).

To quantify the differences between the alleles, we measured the GFP-Sec4 fluorescence intensity of a small corridor adjacent to the PM of the bud and normalized it to the intensity of the whole bud (Fig. 3A). In wild type cells, GFP-Sec4 staining was clearly restricted to the narrow zone beneath the plasma membrane. In *boi1*_Δ_*_86-178_* Δ*boi2*-cells GFP-Sec4 was more equally distributed throughout the bud. GFP-Sec4 was also slightly enriched at the bud neck of these cells (Fig. 3A).

**Figure 3.**
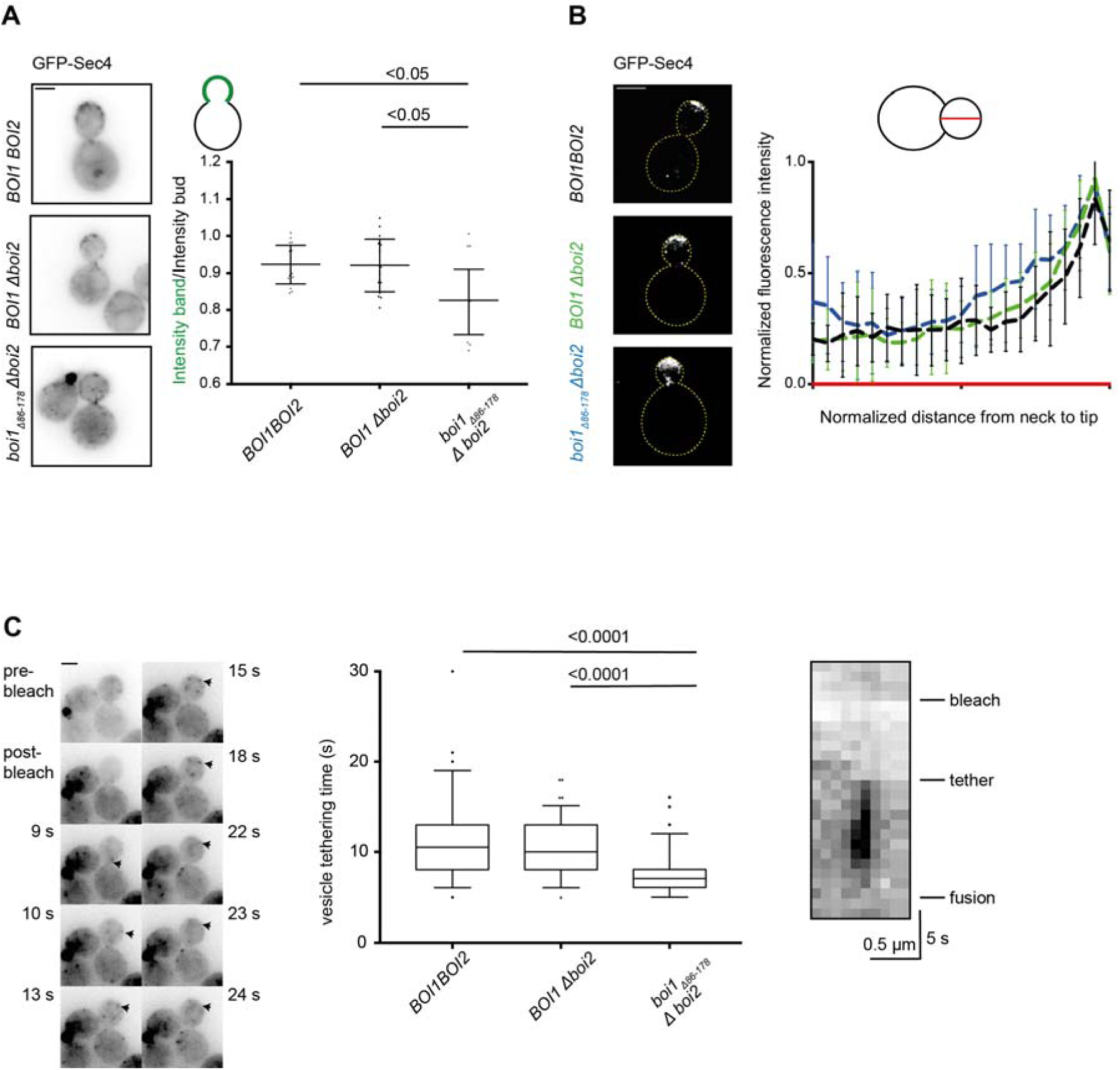
The Boi1-Bud6 interaction is important for vesicle movement and tethering. (A) *BOI1 BOI2-, BOI1* Δ*boi2-*, and *boi1*_Δ_*_86-178_* Δ*boi2*-cells expressing GFP-Sec4 were analyzed by time-lapse microscopy taking an image of 5 stacks every 1s over 104 frames. The bud was bleached after 4 frames to visualize incoming vesicles. Upper panel: Projections of the vesicle distribution of the complete time course. Lower panel: Ratios of mean intensities of a corridor below the plasma membrane representing the tethered and docked vesicles to the mean intensity of the entire bud. Error bars indicate the s.d. of the mean (n=15 from two independent experiments for all genotypes tested). Statistical analysis was performed with a Kruskal-Wallis test followed by Dunńs post-test. (B) *BOI1 BOI2-, BOI1* Δ*boi2-*, and boi1_Δ86-178_ Δboi2-cells expressing GFP-Sec4 were fixed and visualized by SIM. Upper panel: GFP-Sec4 distributions in cells of the indicated genotypes. Mean intensity profiles of GFP-Sec4 from the bud necks to the tips of 10 cells per genotype (Black: *BOI1BOI2*, Green:*BOI1* Δ*boi2*, Blue: *boi1*_Δ_*_86-178_* Δ*boi2*; 5 cells/measurement). Error bars: s.d of the mean. C: Determination of the vesicle tethering time based on time-lapse microscopy from (A). Left panel: Fluorescence microscopy of a wild type cell expressing GFP-Sec4 showing the movement, tethering and fusion of a vesicle in consecutive images. Middle panel: Quantification of vesicle tethering time based on two independent experiments with n*_BOI1, BOI2_*= 102, n*_BOI1,_* _Δ_*_boi2_*= 98, n*_boi1_*_Δ_*_86-178,_* _Δ_*_boi2_*= 96 vesicles measured in total. Statistical analysis is based on a Kruskal Wallis test followed by Dunńs post-test. Right panel: Kymograph used for the determination of the tethering time of the vesicle highlighted in the left panel. Scale bar of all images shown indicates 2 µm.

Tracking individual post-Golgi vesicles is best achieved in medium-sized and large buds. To complement our tracking experiments, we observed GFP-Sec4 by SIM of fixed cells to look at the distribution of post-Golgi vesicles in small buds where vesicular traffic is more tip-directed. Vesicles stained the cortex in a very restricted zone at the tip of wild type cells (Fig. 3B). This zone became slightly broader in *BOI1*Δ*boi2*-cells (Fig. 3B). This trend continued in *boi1*_Δ_*_86-178_* Δ*boi2*-cells where the GFP-Sec4-staining also extended more toward the center of the bud and additionally appeared as small cluster at the bud neck (Fig. 3B).

Vesicles that reached their final destination at the cortex stayed there in average for 10.85 s before disappearing, most probably through fusion with the plasma membrane. The tethering time was significantly reduced to 7.7 s in *boi1*_Δ_*_86-178_* Δ*boi2*-cells. As Δ*boi2*-cells displayed a near wild type tethering time of 10.5 s, we conclude that a loss of the Bud6-interaction site in *boi1*_Δ_*_86-178_* causes a faster fusion of vesicles with the plasma membrane (Fig. 3C).

### Boi1/2 form autonomous actin nucleation sites

The Boi1/2-independent location of Bud6, as well as the prominent presence of heterogeneous, Boi1/2-independent actin structures in the bud prevents the straightforward conclusion that Boi1/2 initiate actin filaments *de novo* (Fig. S3). By fusing Boi1 to Pex3, a membrane protein of the peroxisomes, we removed Boi1 from the known actin nucleation centers of the bud to study its activity in isolation at the membrane of the peroxisome (Fig. 4A) (Luo, Zhang, Guo 2014). Pex3_1-45_mCherry-Boi1 (Pex-Boi1) is efficiently targeted to the membrane of the peroxisomes (Fig. 4B). By co-expressing a GFP fusion to proteins involved in polar growth including known ligands of Boi1/2, we could show that Boi1 attracts all its tested binding partners to the peroxisome including members of the exocyst (Fig. 4C, 4E, S4).

**Figure 4.**
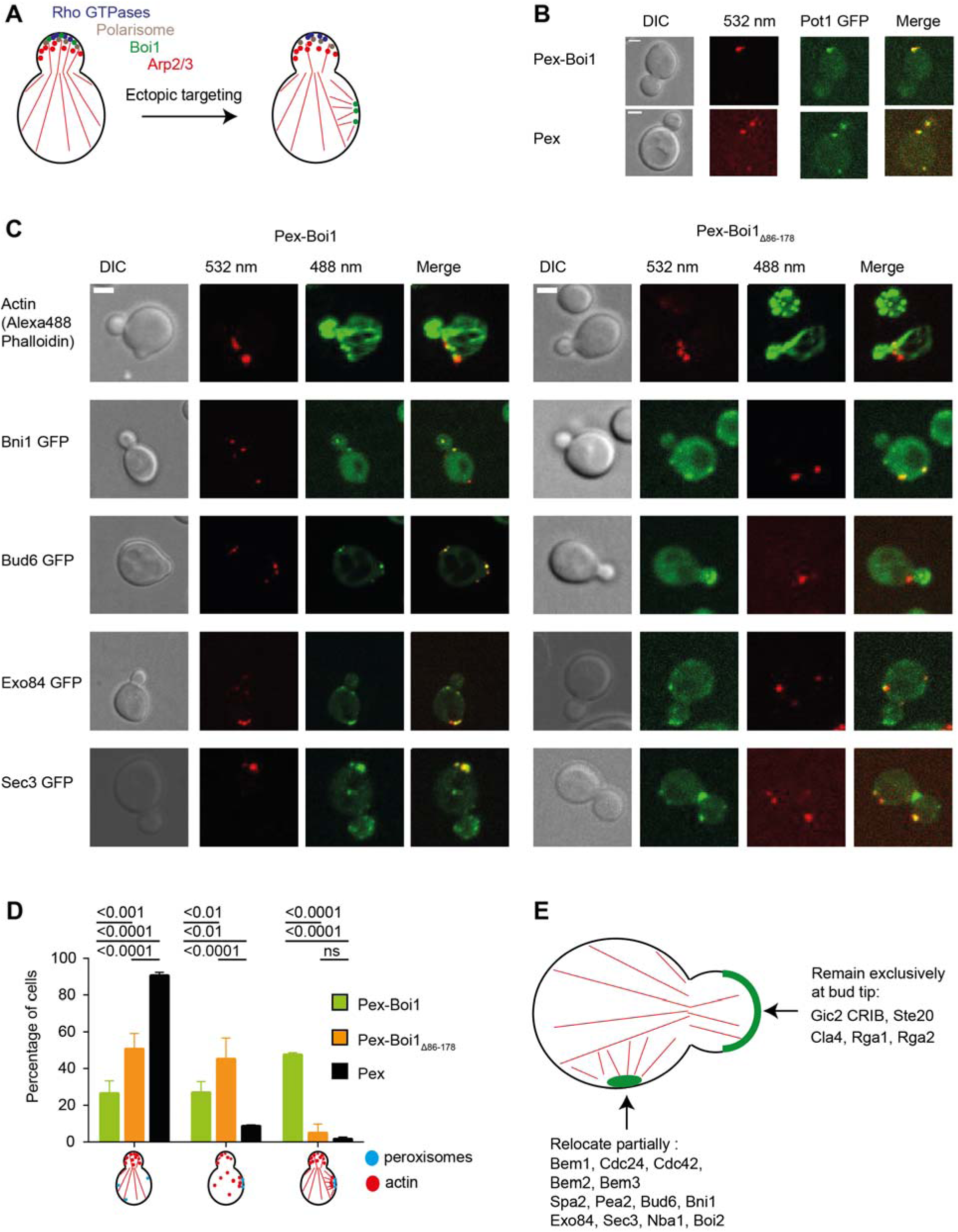
A Boi1-Bud6-Bni1 complex initiates actin filaments *de novo*. (A) Experimental design: Pex3_1-45_-mCherry was fused to Boi1 (Pex-Boi1) or to Boi1_Δ86-178_ (Pex-Boi1_Δ86-178_) to target the fusion to the peroxisome allowing to study its actin nucleation activity in isolation from other nucleation factors in the bud. (B) Cells co-expressing the peroxisomal marker Pot1-GFP and Pex-Boi1 were visualized by confocal microscopy. Representative images show maximum projections of 10 z stacks and a clear co-localization of both fusion proteins. (C). Fluorescence microscopy of cells co-expressing Pex-Boi1, or Pex-Boi1_Δ86-178_ together with GFP-labeled Bni1, Bud6, Exo84, or Sec3. Phallodin-Alexa488 staining was used to visualize the actin cytoskeleton of these cells. Scale bars correspond to 2 µm. (D) Phenotypic quantification of the actin cytoskeleton of cells expressing the indicated color-coded Pex-fusions after Alexa488-Phalloidin staining. The categorization into three phenotypes is shown in the cartoons: I) Polarized actin cables and patches; II) Mis-localized actin patches, partially co-localizing with peroxisomes; III) Bi-polar actin cable organization with filaments arising from peroxisomes. Data were derived from three independent actin stainings with in total n_Pex3-Boi1_= 553, n_Pex3-mcs_= 598, n _Pex3-Boi1Δ86-178_= 650 cells. Bars indicate the mean, and error bars the s.d.. 2-way ANOVA followed by a Tukeýs Multiple comparison test were used to determine the significance. (E) Summary of the distributions of different GFP fusions in Pex-Boi1 expressing cells. A quantitative analysis of the distributions is shown in Table S2.

Actin staining revealed the establishment of an alternative axis of cell polarity in the Pex-Boi1-expressing cells (Figs. 4 C, D). The actin cables of this alternative axis seem to emanate from peroxisomes located in the mother. To distinguish the contribution of the actin nucleation factors from the contributions of all other recruited proteins, we repeated the experiments with cells expressing peroxisome-targeted Boi1_Δ86-178_ (Pex-Boi1_Δ86-178_) lacking the binding site to Bud6. Accordingly, Bud6-GFP was not longer found at Pex-Boi1_Δ86-178_-labelled peroxisomes, whereas Bni1, the exocyst subunit Exo84 as well as other members of the polarity complex still co-localized (Figs. 4C, S4, Table S2). Actin cables in this strain were often less polarized towards the cell tip but did not any longer align towards the peroxisomes (Fig. 4C, D). A significant portion of cells still contained actin patches around peroxisomes (Fig. 4D).

The GFP fusion of the v-SNARE Snc1 (GFP-Snc1) and Sec4 were also enriched at the peroxisomes of Pex-Boi1-but not of Pex-Boi1_Δ86-178_ expressing cells (Fig. 5A, B). As both GFP-fusions are attached to post-Golgi vesicles, their recruitment to the peroxisome indicates the reconstitution of an at least partially functional vesicle-tethering zone. Consequently, Boi1-but not Boi1_Δ86-178_-labelled peroxisomes, when found in close apposition to the PM, often induce an outward bulging of the cell wall (Figs. 4C, 5A).

**Figure 5.**
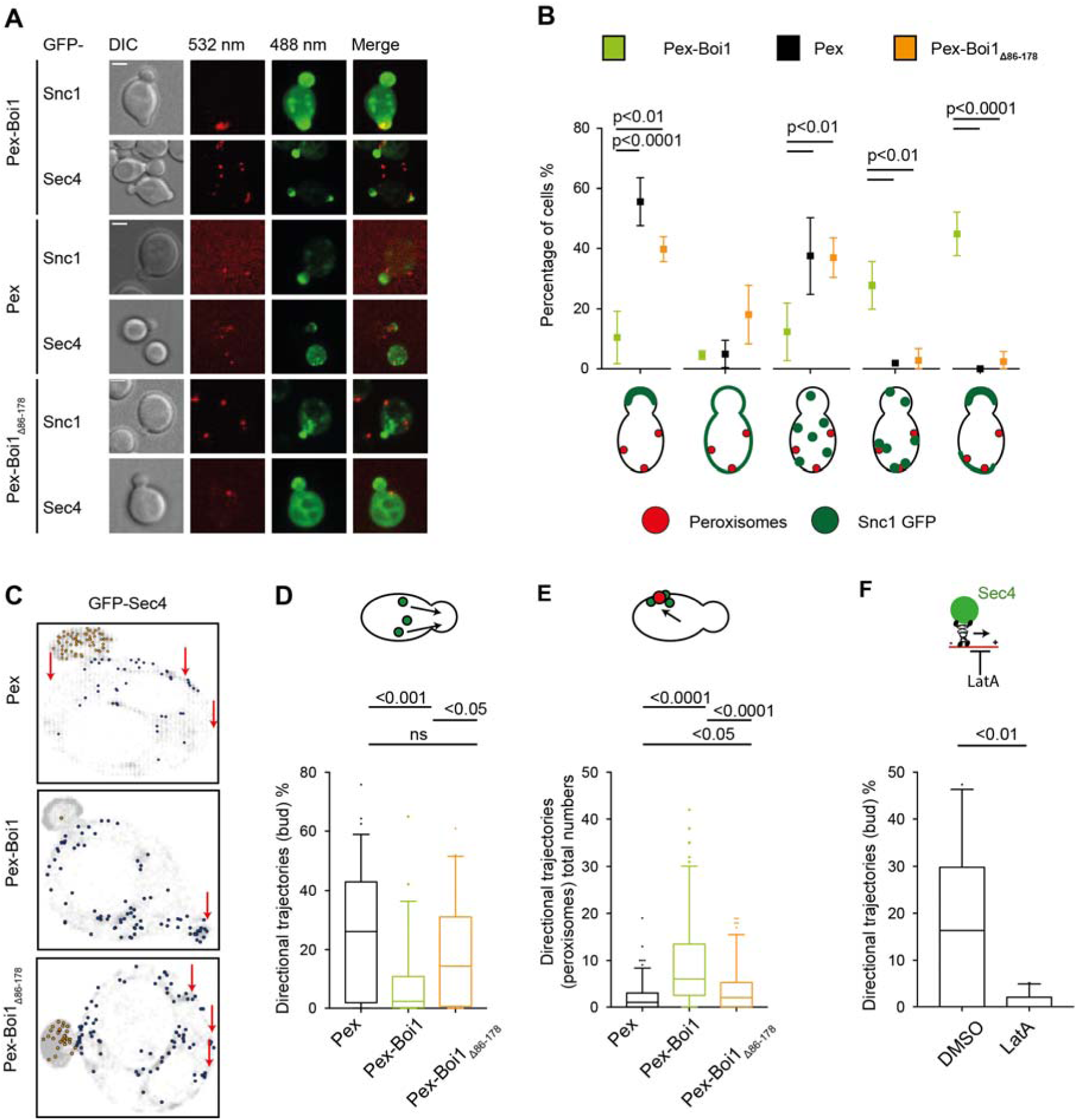
Pex-Boi1 redirects the flow of post-Golgi vesicles to peroxisomes. (A) Representative images of cells co-expressing GFP-Sec4, or the v-SNARE GFP-Snc1, and either Pex-Boi1, Pex-Boi1_Δ86-178_, or Pex. Scale bar indicates 2 µm. (B) Quantification of GFP-Snc1 distributions in cells of (A) (Pex-Boi1 (green), Pex-Boi1_Δ86-178_ (orange), Pex (black)). Distributions were categorized according to the cartoons in the lower panel. The experiments were performed twice with in total n_Pex3-Boi1_= 177, n_Pex3-mcs_= 107, n _Pex3-Boi1Δ86-178_= 221 cells. Statistical analysis was performed with a two-way-ANOVA and Tukeys-multiple comparison post-test. Error bars indicate s.d. of the mean. (C) Δ*boi1* cells co-expressing Pex-mCherry, Pex-Boi1, or Pex-Boi1_Δ86-178_ together with GFP-Sec4 were visualized by confocal microscopy. After bleaching the bud, images were taken every 100 ms over 35 s to follow the movement of the labeled vesicles. Shown are single cells of each genotype displaying the destinations of all vesicles. Vesicles moving into the bud are labeled orange, vesicles showing directional movement in the mother are painted blue, and the remaining vesicles displaying no, or random movements are in grey. Red arrows indicate the position of the peroxisomes. (D) Quantification of vesicle trajectories of cells shown in (A). Left panel: fraction of trajectories of vesicles moving into the bud of cells expressing the indicated Pex3 fusions. Right panel: Number of directional trajectories/cell of vesicles moving towards the peroxisomes displaying the indicated Pex3 fusions. The percentage of vesicles showing directed movement into the bud was calculated in all three alleles shown in (B). Black: Δ*boi1* cells expressing Pex3_1-45_mCherry (n=69 cells); Green: Δ*boi1* cells expressing Pex-Boi1 (n=53 cells); Orange: Δ*boi1* cells expressing Pex-Boi1_Δ86-178_ (n=44 cells). Data were collected from three independent measurements. Statistical analysis was performed with a Kruskal-Wallis test followed by Dunńs post-test. (E) Directed movements of the vesicles are strictly actin-dependent. Cells co-expressing Pex3_1-45_mCherry and GFP-Sec4 were treated with 100 µM LatA or DMSO (control). Images of a single plane were taken every 100 ms over 35 s. The percentages of directionally transported vesicles entering the bud were calculated using 24 DMSO-treated, and 22 LatA-treated cells from two independent experiments. Statistical comparison was performed with a Mann-Whitney test. Boxplots in (D) and (E) display the median and 5 to 95 percentile whiskers.

Tracking of individual GFP-Sec4-labeled post-Golgi vesicles confirmed the formation of an alternative and functional polarity axis (Video_S2) (Fig. 5C, D). Cells containing Boi1-decorated peroxisomes displayed a reduced flux of vesicles to the bud (Fig. 5C, D). The reduction was accompanied by an increase in the fraction of vesicles that moved away from the bud towards the Boi1-labed peroxisomes (Fig. 5C, D). This redirection of vesicular traffic was not seen in cells expressing Pex-Boi1_Δ86-178_ (Fig. 5C, D). The measured directional vesicular traffic was strictly actin-dependent (Fig. 5E).

## Discussion

In most eukaryotic organisms post-Golgi vesicles arrive at the cell plasma membrane through a directed long-distance walk on microtubules or actin cables. The vesicles are then handed over to the actin filaments underlying the cortex (Hume et al. 2011; Porat-Shliom et al. 2013). The actin-bound vesicles are either kept on hold during regulated exocytosis or processed directly for docking and fusion. In budding yeast post-Golgi vesicles are transported exclusively on actin cables to the plasma membrane of the bud (Pruyne, Schott, Bretscher 1998). We propose that in budding yeast, similar to other eukaryotes, secretory vesicles switch from actin cables used for long-distance transport to cortical actin filaments that guide the vesicle to the docking and fusion zone. Our experiments point to Boi1/2 as nucleation sites for actin filaments below the plasma membrane. Boi1/2 locate at the cortex and bind to Bud6 and the formin Bni1, two proteins that form a potent actin nucleation and elongation complex (Graziano et al. 2011; Graziano et al. 2013; Moseley and Goode 2005). Abrogating the interaction between Boi1/2 and Bud6 reduces actin cable density in buds, increase the random movement of vesicles, and shortens the tethering time of the bound vesicles. Furthermore, the artificial relocation of Boi1 to peroxisomes creates an alternative tethering zone at the peroxisome including secretory vesicles and actin filaments that emanate from these sites. The zone is formed by the many binding partners of Boi1 and depends on Boi1’s ability to recruit Bud6 to initiate actin cables (Fig. 6). The efficacy by which the peroxisome-tethered Boi1 competes with other factors in the cell for actin filament formation might suggest that Boi1/2 not only anchors but also activates the Bud6/Bni1 complex upon binding.

**Figure 6.**
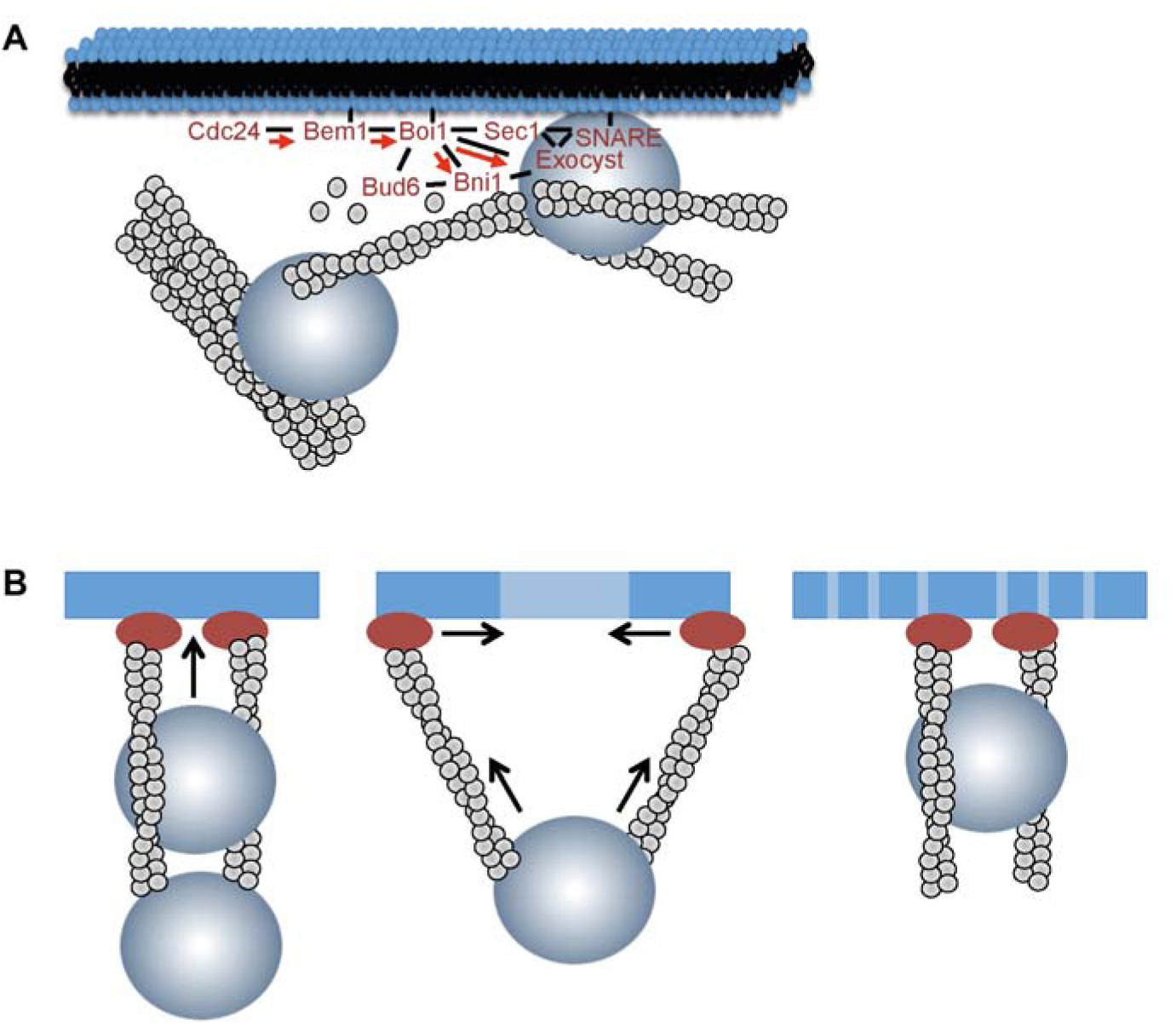
The Boi-proteins as hubs of polarized secretion and actin filament nucleation. (A) Boi1 binds through its PH domain to the plasma membrane of the bud and anchors together with phospho-lipids the Bem1-Cdc24 complex to the membrane. Cdc24 activates Cdc42 that is shuttled (red arrows) through Bem1 and Boi1 to its effectors. The Boi1-bound Bud6-Bni1 complex initiates and elongates filaments. Those serve as tracks for post-Golgi vesicles to the membrane where they are fixed and kept from spontaneous fusion. In addition Boi1 contacts the exocyst and the SM protein Sec1 to prepare the vesicle for SNARE-mediated fusion (Kustermann et al. 2017). (B) The Boi1/2- containing receptor complexes generate the actin filaments that serve as tracks for the incoming vesicles. Upon fusion, the membrane of the vesicle pushes the receptor complexes apart. The Myo2-driven movement of the next vesicle contracts the receptors to their original position, thus providing directional persistency of the vesicle fusion process.

By focusing our analysis on the *boi1*_Δ_*_86-178_*-allele, we tried to separate its influence on actin filament formation from the protein’s two other main functions, the localization of the Bem1-Cdc24 complex and the formation of the tethering and docking complex (Bender et al. 1996; Kustermann et al. 2017). The former activity is located on a short binding motif in the middle of the sequence of Boi1, whereas the latter activity locates on the membrane- and Sec1-binding C-terminal PH domain, and the Exo84-, Sec3-binding N-terminal SH3 domain (Bender et al. 1996; Kustermann et al. 2017). We propose that the concentration of all three activities in one protein coordinates vesicle fusion with trafficking and enables the control of both activities through Rho-GTPases (Fig. 6A). Linking vesicle tethering/fusion with actin nucleation might foster vesicle docking at sites where fusion has recently occurred and thus equip secretion with the processivity that is required for polarized growth (Fig. 6B). Binding to Bem1-Cdc24 might channel the activated Cdc42 through the Cdc42_GTP_-binding PH domain of Boi1/2 directly to Bni1, and the exocyst components Sec3 and Exo70 that tether the vesicle to the membrane and together with the other exocyst components catalyze the formation of the docking complex (Fig. 6A) (Adamo et al. 2001; Guo et al. 1999; Guo, Tamanoi, Novick 2001; He et al. 2007; Morgera et al. 2012; Yue et al. 2017).

The reduction in vesicle tethering time through the dissolution of the Boi1-Bud6 complex seems counterintuitive, yet might indicate that the cortical actin in yeast as in higher eukaryotes not only directs movement to the membrane but also restricts and controls the access of the vesicles to the plasma membrane (Li, P. et al. 2018; Meunier and Gutierrez 2016). This conclusion is supported by the phenotype of cells carrying a *bud6* deletion. Here the number of actin cables is reduced, but the velocity of post-Golgi vesicles, and the randomness of their movements is increased. At the same time exocytosis became less efficient in these cells (Jose et al. 2015).

Although budding yeast uses only actin structures for transport and docking, the significance of our findings is not restricted to these cells. Studies in neuroendocrine cells showed that the role of the cortical actin cytoskeleton is quite similar with respect to the coordination of exocytosis, where it also directs vesicular flow, and mediates docking and fusion (Chasserot-Golaz et al. 2005; Gabel et al. 2015; Gonzalez-Jamett et al. 2017; Maucort et al. 2014). Upon stimulation, actin associated proteins like the actin bundling protein AnnexinA2 are targeted to the SNARE complex at the plasma membrane to reorganize the integrity of the cortical actin cytoskeleton and generate a vesicle fusion-promoting environment (Gabel et al. 2015; Umbrecht-Jenck et al. 2010). The fission yeast *S.pombe* transports post-Golgi vesicles on microtubules. Deletion of its single Boi homologue Pop1 leads to the accumulation of secretory vesicles in the cytosol (Nakano et al. 2011). Pop1 was also shown to bind to the *S.Pombe* formin For1. Disrupting this interaction disturbs the actin cytoskeleton (Rincon et al. 2009). Pop1 complements the essential function of Boi1/2 in vesicle fusion (Kustermann et al. 2017). These findings indicate that the molecules and mechanisms involved in the transfer of secretory vesicles from their long-distance carrier to cortical actin structures are quite conserved.

## Materials and Methods

### Growth conditions and cultivation of strains

All yeast strains in this study are derivatives of the *Saccharomyces cerevisiae* JD47 strain. Cells were incubated at 30°C in YPD or synthetic medium lacking specific amino acids or complemented with antibiotics for selection. *E.coli* XL1 blue cells were used for plasmid amplification and grown at 37°C in LB medium containing antibiotics. *E.coli* BL21 cells were used for protein production and were grown in LB or SB medium at 37°C or 18°C.

### Construction of plasmids and strains

Detailed lists of all primers and plasmids from this study are provided in Tables S3 and S4. Fusions of GFP or CRU to *BUD6*, *BNI1*, *BOI1* or *boi1*_Δ_*_86-178_* were constructed by PCR amplification from genomic DNA of the respective C-terminal ORFs without stop codon as described (Dunkler, Muller, Johnsson 2012; Fujimura-Kamada, Hirai, Tanaka 2012; Kustermann et al. 2017; Manavalan and Johnson 1987; Norden et al. 2006; Wittke et al. 1999). The obtained DNA fragments were cloned via *Eag*I and *Sal*I restriction sites in front of the CRU-, GFP-, mCherry-module on a pRS303, pRS304 or pRS306 vector (Sikorski and Hieter 1989). For integration into the genome, the plasmids were linearized using a single restriction site within the C-terminal genomic DNA sequence. Successful integration was verified by PCR of single yeast colonies with diagnostic primer combinations using a forward primer annealing in the target ORF but upstream of the linearization site, and a reverse primer annealing in the C-terminal module. Gene deletions were obtained by replacing the ORF with an antibiotic resistance cassette through single step homologous recombination as described (Janke et al. 2004). Genomic N_ub_ fusions were obtained as described (Hruby et al. 2011). Generation of yeast centromeric plasmids containing N_ub_ fusion proteins included initial PCR amplification of indicated fragments from genomic DNA containing a SalI restriction site in the forward- and Acc65I restriction site in the reverse primer and ligated into the plasmid *N_ub_-empty kanMX4*.

Fragments of *BUD6*, *BNI1* or *BOI1* were expressed as GST- or 6xHis-fusions in *E.coli* BL21. GST-fusions were obtained by amplification of the respective fragments from genomic yeast DNA using primers containing NcoI/EcoRI restriction sites. The PCR fragments were cloned in-frame behind GST in the plasmid pGex6P1 or pGex2T (GE Healthcare, Buckinghamshire, UK). For 6xHis-tagged fragments, a PCR of the respective fragment from genomic DNA using primers containing *Sfi*I restriction sites was performed and the product was inserted in frame downstream of a 6xHis-tag into the pAC plasmid (Schneider et al. 2013). The chimeric Pex3_1-45_mCherry pRS306 plasmid was adapted from Luo et al. (Luo, Zhang, Guo 2014). *BOI1* or *boi1*_Δ_*_86-178_* were amplified from genomic DNA and inserted in frame behind the mCherry tag using BamHI/SalI restriction sites.

Genomic integration of the *boi1*_Δ_*_86-178_* allele was performed by „delitto perfetto“ methodology or CRISPR CAS9 (Laughery et al. 2015; Storici and Resnick 2006). The successful deletion/exchange of amino acids were confirmed by sequencing of single-colony PCRs. A detailed description of the construction of all plasmids can be obtained upon request.

### In vitro binding assays

#### Protein expression

O.n. cultures of *E.coli* BL21 cells were diluted to OD_600_= 0.3, and incubated at 37°C in LB or SB medium to an OD_600_= 0.8 before protein synthesis was induced by the addition of IPTG. Protein expression conditions were optimized for each expression construct (see Table S5). Cell pellets were stored after induction at −80°C.

#### Cell extract preparation

Cell pellets were resuspended in 1x PBS or 1x HBSEP (pH 7.4, 10 mM Hepes, 150 mM NaCl, 3 mM EDTA, 0.005% Tween-20) containing 1x protease inhibitor cocktail (Roche Diagnostics, Penzberg, Germany), incubated for 20 min with 1 mg/ml lysozyme on ice, and subsequently subjected to sonification for 2×4 min with a Bandelin Sonapuls HD 2070 (Reichmann Industrieservice, Hagen, Germany). Lysates were spun down at 40.000g for 10 min/4°C. Supernatants were transferred either directly to the binding assay, or used for further purification.

#### Binding assay

All incubation steps were carried out under rotation in the cold room. Extracts of GST or GST-fusion proteins were incubated for 0.5-1h with glutathione-coated sepharose beads (GE Healthcare, Freiburg, Germany) equilibrated in PBS (Boi1-Bni1) or HBSEP (Boi1-Bud6). Beads were washed twice and incubated with 0.1 mg/ml BSA (Boi1-Bni1) (Sigma Chemicals, St.Louis, USA) for 30 min before the beads were treated with either purified 2 µM 6xHis Bud6_1-141_, or extract of 6xHis Boi1_1-300_ in the presence of 0.1 mg/ml BSA for 1h. Beads were washed 3 x with HBSEP or PBS before eluting the bound protein with 1xGST elution buffer (pH 8.0, 50 mM Tris, 20 mM reduced glutathione). Protein eluates were separated by SDS PAGE and stained with Coomassie Brilliant Blue or with anti-His antibody after transfer onto a nitrocellulose membrane (Sigma-Aldrich, Steinheim, Germany; dilution 1: 5000).

#### Quantification of Western blots

Western blots were quantified with ImageJ. A detailed step-by-step procedure is given in (University of Queensland, Diamantina Institute, 27 April 2017, https://di.uq.edu.au/community-and-alumni/sparq-ed/sparq-ed-services/using-imagej-quantify-blots). Briefly, the histogram of the intensities of each band of a western blot was used to calculate the area under the curve (AUC), which correlates to the size and brightness of each band. The AUC of each band was normalized to the AUC of the input band to quantitatively compare the amount of bound 6xHis-tagged fusion protein in each lane of the gel.

#### Protein purification

For purification of 6xHis-Bud6_1-141_ cell pellets were extracted as above in 1x IMAC binding buffer (pH 7.5, 300 mM NaCl, 50 mM KH2PO4, 20 mM Imidazole). Purification was achieved by immobilized metal affinity purification followed by size exclusion chromatography (SEC) on an ÄktaPurifier Chromatography System (GE Healthcare, Buckinghamshire, UK).

The final protein concentration was determined with a NanoDrop ND-1000 spectral photometer (Peqlab, Erlangen, Germany) at 280 nm excitation, and based on a calculated excitation coefficient of 8.5 mM^-1^cm^-1^ and a molecular weight of 17.7 kDa (www.expasy.org). The purified protein was used either directly for pull down analysis or stored at −20°C.

### In vivo interaction analysis with the Split-Ubiquitin system

Split-Ubiquitin array analysis: A library of 533 different α-strains each expressing a different N_ub_ fusion were mated with a Bni1CRU-, Bud6CRU-, Boi1CRU- or Boi1_Δ86-178_CRU-expressing a-strain. Diploid were transferred as independent quadruplets on SD media containing 1 mg/ml 5-FOA, and different concentrations of copper to adjust the expression of the N_ub_ fusions (Fig. S1) (Dunkler, Muller, Johnsson 2012). Individual Split-Ub interaction analysis: CRU- and Nub-expressing strains were mated or co-expressed in haploid cells, and spotted onto medium containing 1 mg/ml FOA and different concentrations of copper in four 10-fold serial dilutions starting from OD600=1. Growth at 30°C was recorded every day for 2 to 5 days.

### Microscopy

#### Wide-field and confocal microscopy

Fluorescence microscopy was performed on an Axio Observer Z.1 spinning-disc confocal microscope (Zeiss, Göttingen, Germany) containing a switchable Evolve512 EMCCD (Photometrics, Tucson, USA), or an Axiocam Mrm camera (Zeiss, Göttingen, Germany). The microscope was also equipped with a Plan-Apochromat 100×/1.4 oil DIC objective, and 488 nm, 561 nm, and 635 nm diode lasers (Zeiss, Göttingen, Germany). Images were recorded with the Zen2 software (Zeiss, Göttingen, Germany), and analyzed with FIJI. Alternatively, time-lapse microscopy was performed with a DeltaVision system (GE Healthcare, Freiburg, Germany) provided with an Olympus IX71 wide field microscope (Olympus, Hamburg, Germany). This microscope contained a CoolSNAP HQ2-ICX285, or a Cascade II 512 EMCCD camera (Photometrics, Tucson, USA), a 100× UPlanSApo 100×1.4 Oil ∞/0.17/FN26.5 objective (Olympus, Münster, Germany), a steady-state heating chamber, and a Photofluor LM-75 halogen lamp (89 NORTH, ChromaTechnology, Williston, USA).

Secretory vesicles in the bud tip were observed with an iMIC microscope (Photonics, Pitsfield, USA) equipped with an Andor-Clara camera (Type Clara DR 328G-C01-SIL CCD), an Oligochrome (Ex 390/40; 482/20; 563/20; 640/20), a LED camera with excitation filters FF01-340/26-25, FF01-387/11-25, and FF01-470/22-25, and a 60xOil Olympus ApoN 60x/1.490oil∞/0.13-0.19 objective. The microscope was controlled with the software Live Acquisition (v.2.6.0.34) (FEI Munich, Gräfelfing, Germany). Images were taken in 5 z sections (Δz= 0.5 µm) with an excitation time of 80ms and a laser intensity of 90%.

#### Structured illumination microscopy (SIM)

Imaging setup is described in Rüthnick et al. (Rüthnick et al. 2017). Shortly, Alexa488-Phalloidin stained cells were suspended in PBS and imaged with a Nikon N-SIM system equipped with total internal reflection fluorescence Apochromat 100× 1.49 NA oil immersion objective, and a single photon–detection electron-multiplying charge-coupled device camera (iXon3 DU-897E; Andor Technology, Belfast, UK) using a 488 nm Laser for excitation with an emission bandpass filter 520/45. Reconstructions were performed with the image analysis software (Nikon, Tokio, Japan). Images were taken in 5 z sections.

#### Quantitative analysis of fluorescence microscopy

All microscopy files were processed and analyzed with FIJI (US National Institute of Health; Version 2.0.0-rc-69) (Schindelin et al. 2012). Images were acquired as 5 to 14 z-stacks and analyzed either in single layers or as projections of single layers.

The number of actin filaments running in parallel to the mother-bud axis within the bud or mother of yeast cells was determined with the assistance of the FIJI tool “Plot profile”. Filaments were counted at a position half way in the bud or half way in the mother compartment. Local intensity maxima were considered as actin filament counts. Quantification in the local enrichment of fluorescence intensity (RI= Relate Intensity) was calculated base on the formula:

RI=(I ROI-IBackground)/(ICytosol-IBackground). IROI describes the mean intensity within the region of interest (ROI), IBackground describes the mean intensity within a region outside of the cell, and ICytosol describes the mean intensity within a region of the cytosol of the mother compartment.

To determine the local enrichment of GFP fusion proteins at peroxisomes, the mean intensity in the GFP channel at all mother cell-located peroxisomes was averaged to determine IROI. The coefficient of variation (COV) used to determine actin cable staining density was calculated as the ratio of standard deviation (s.d.) to the mean fluorescence intensity of the whole mother cell or bud (Garabedian et al. 2018). A less dense actin cable network contains larger dark, cable-free regions. This results in a higher s.d. and a lower mean fluorescence intensity, thus increasing the COV.

The vesicle distribution in the bud of cells expressing GFP-Sec4 was determined from SIM images with the FIJI tool “Plot profile” by setting a ruler from the bud neck to the bud tip. The relative length was divided into 5% steps from 0% at the bud neck to 100 % at the tip. Along this line (thickness: 0.16 µm = 5 Pixel) the mean fluorescence intensity was determined and the background subtracted. The mean intensities along this line represent relative intensities normalized to the highest mean intensity. 10 cells with bud lengths between 2 and 2.5 µm were analyzed for each genotype.

#### SIM image processing

Images were stacked to maximum projections. Background subtraction for the determination of actin cable numbers and actin cable staining density in bud and mother was performed with the “running ball” method in FIJI with a radius of 20 pixels (=0.64 µm). To subtract the background for determination of the vesicle distribution in the bud, the highest intensity value in a vesicle-free cytosolic area of the mother was subtracted from all intensities in the bud.

### Actin staining

Exponentially-grown cells were fixed for 10 min by adding 3.7% Formaldehyde to the medium. Cells were resuspended in 3.7% Formaldehyde (in 100 mM KH_2_PO_4_), and incubated for 1h before buffer was exchanged to 1 µM ethanolamine (in 100 mM KH_2_PO_4_) for 10 min. Cells were washed twice with PBS, and incubated with 66 nM Alexa-fluorophore-conjugated phalloidin (Thermo Fisher Scientific, Waltham, MA, USA) for 30 min or over night at 4°C. Actin patches were removed by addition of 100 µM CK-666 (Merck, Darmstadt, Germany) to the cell medium at 30°C, 10 min before cells were fixed. Samples that were imaged by SIM were prepared with buffers that were filtrated with a 0.22 µm syringe filter (TPP, #99722, Trasadingen, Switzerland) prior to usage.

### Vesicle tethering analysis

Exponential growing cells were embedded between a coverslip and a custom-made glass slide (Glassbläserei, Uni Ulm) into a 3.8% agarose gel (containing 1xSD medium). Cells were imaged at room temperature (approx. 22-26 °C). To keep conditions constant among different genotypes, we measured the corresponding cells in a defined order and reversed this order during the repetition. To follow individual vesicles in the bud, images (5 z-stacks, 0.5 µm per stack) were taken every second. Each time-lapse series included 4 images that were acquired before and 100 images acquired after bleaching the bud of the cell. Tethering time was determined as the time interval between the arrival of a vesicle at the cortex and the time when it disappears. Only vesicles were taken into account that stayed fixed for more than 4 s at the cortex and were visible in the middle three layers.

#### Vesicle distribution in the bud

To quantify the vesicle distribution of incoming vesicles in the bud over the whole time course of 100 images, the middle three layers were stacked to maximum projections. Maximum projections of the consecutive 100 images were further maximum projected into a single image. The mean intensity of a corridor below the plasma membrane of the bud was determined using the “segmented line” tool in FIJI with a line width of 3 pixels (= 0.326 µm), and normalized to the mean intensity of the whole bud. Both mean intensity values were background-subtracted before calculating the ratio.

### Vesicle tracking analysis

#### Image Acquisition

Yeast cells expressing GFP-Sec4 from a centromeric plasmid under a methionine adjustable P*_MET17_*-promoter were grown in selective medium lacking methionine. Exponentially grown cells were spun down, and resuspended in selective medium. 3.1 µl of the cell suspension were embedded between a glass microscopy slide and a cover slip. Microscopy was performed at room temperature. Images were taken with a confocal microscope over a time course of 35 s using a single z-stack and acquiring images every 100 ms (Intensity Laser 488nm= 20%; excitation time= 50ms). Before the start of each time lapse, an image of the green and red channel was taken to reconstruct the position of peroxisomes in the trajectory plots for post-analysis. The bud of the cells and vesicle-dense regions (in Pex3-Boi1) in the cortex of the mother were subsequently bleached to increase the signal to noise ratio of single vesicles in the mother and daughter cell.

#### Vesicle tracking

Time-lapse images were analyzed with the MOSAIC suite plugin for Fiji/ImageJ (Sbalzarini and Koumoutsakos 2005; Schindelin et al. 2012). To observe and follow vesicles over time, we adjusted the following parameters, whereas all other variables were left in default mode: For particle detection: i) Particle radius: 0.133 µm; ii) Percentile (r): 0.3-1.5 (value was adjusted and vesicle detection was verified in randomly chosen images of the time lapse by using the „preview detected“ function). For particle linking: i) Link range: 5 images; ii) Dynamics: Brownian; iii) Displacement: 1.33 µm/100 ms.

#### Directionality analysis

Directionality analysis based on microscopy coordinate measurements of trajectories was performed in R statistical package (R Core Team 2013). For each cell, we first fitted a linear function delineating the mother cell and the bud. This function was later used to select trajectories originating from the mother cell for downstream directionality analyses. For each trajectory measured over a time span within a given cell, we calculated two Euclidean distance measurements: (1) distance between two subsequent time points and (2) Absolute Euclidean distance (displacement) from the original position. We then took the ratio of the net displacement to the cumulative distance traveled, with a maximum attainable upper limit value of 1 representing a perfectly directional trajectory moving in a straight line. For ease of interpretation, we took the log of this ratio, setting the maximum value to 0 while all variations of movement would fall in the negative range. In the final step, we measured area under the curve (AUC) of the calculated ratios along the entire length of the trajectory. To test the statistical significance of the directional movement against a random motion, we generated a null hypothesis distribution using a Monte Carlo simulation. To achieve this, coordinate measurements of each trajectory were first randomized and Euclidean and AUC measurements were calculated by following the steps discussed above. This was iterated 1000 times for each trajectory to finally generate the null distribution. Significance test of the empirical AUC was performed using the formula *p*= (*r* + 1)/(*N* + 1), where N is the total number of iterations (1000), r counts the number of events in the simulation model that yielded AUC values greater than the empirical/observed AUC, and p is the calculated p-value (Davison and Hinkley 1997). The decision on whether a given trajectory was moving either towards or away from the bud was made strictly based on its final destination as determined by the linear function delimiting the mother cell and the bud. For final percentage of directional trajectories calculations, we excluded short trajectories with lengths less or equal to 9. Statistical comparisons of directionality measurements among genotypes was performed using the non-parametric Kruskal-Wallis one-way ANOVA followed by Dunn’s pairwise post-hoc test.

#### Movement towards peroxisomes

To quantify the number of vesicles moving towards peroxisomes, we overlaid a plot containing vesicle destinations (generated with R) with an image of both fluorescent channels taken right before time-lapse microscopy. Peroxisomes at the mother cortex are mainly immobile, allowing us to combine still images with subsequent time-lapse analysis (Fagarasanu et al., 2006). The directionality plot contained i) the destinations of vesicle trajectories that were calculated to move directional but away from the bud (not ending in the bud) and ii) the destination of all trajectories. The latter were required to overlay and correctly fit the proportions of the raw microcopy image with the directionality plot in order to locate the peroxisomes on each plot. We counted the number of trajectories moving away from the bud that co-localized or directly neighbored peroxisomal sites within each cell of a given allele.

## Author contribution

O. Glomb, L. Rieger, Y. Wu, and N. Johnsson designed and analyzed the experiments. D. Rüthnick and O. Glomb performed and analyzed the SIM experiments. Y. Wu, L. Rieger and O. Glomb performed the protein interaction screens and analyses. O. Glomb analyzed vesicular traffic and together with Y. Wu the actin cytoskeleton. O. Glomb investigated the Pex3-Boi1 expressing cells. M. Mulaw wrote the software for vesicle analysis and statistically evaluated together with O. Glomb the tracking experiments. N. Johnsson and O. Glomb designed the study. O. Glomb and N. Johnsson wrote the manuscript.

## Acknowledgement

We thank the Nikon Imaging Facility of Heidelberg for granting access to SIM and its analysis. We thank the Institute for General Physiology at Ulm University for providing introduction and access to the IMIC microscope. We thank Ute Nussbaumer, Steffi Timmermann, and Nicole Schmid for strain construction, and Drs. Judith Müller, Alexander Dünkler and Reinhild Rösler for sharing constructs, advice and strains. The work was funded by grants from the DFG to N.J. (Jo 187/5-2; Jo 187/8-1). The authors declare no conflicts of interest.

