## Supplementay Figures and Tables for "An actin nucleation complex catalyzes filament formation at sites of exocytosis"

### Supplementary Figures

#### Figure S1

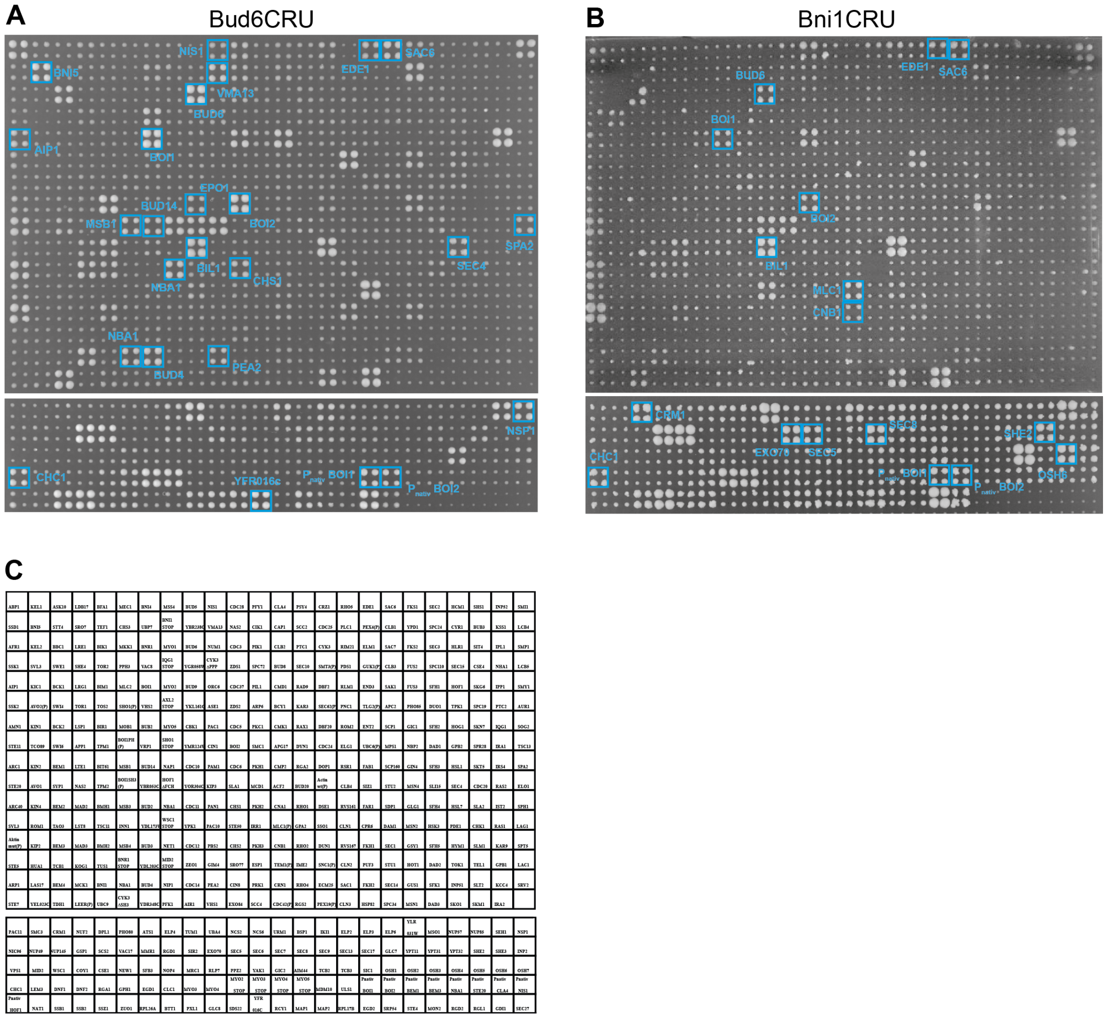


Determination of Bud6 and Bni1 interaction partners *in vivo*. Split-Ub interaction assays of 504 yeast strains co-expressing Bud6CRU (A), or Bni1CRU (B), each with a different $N_{ub}$ fusion protein. Cells were mated and spotted in quadruplets. Shown are the growths of the yeast strains after two (A) or four (B) days on medium containing 5-FOA. Positive hits are indicated by blue frames. Remaining hits are either false positives or interactions by $N_{ub}$-labeled chaperones(Hruby et al. 2011).

### Figure S2

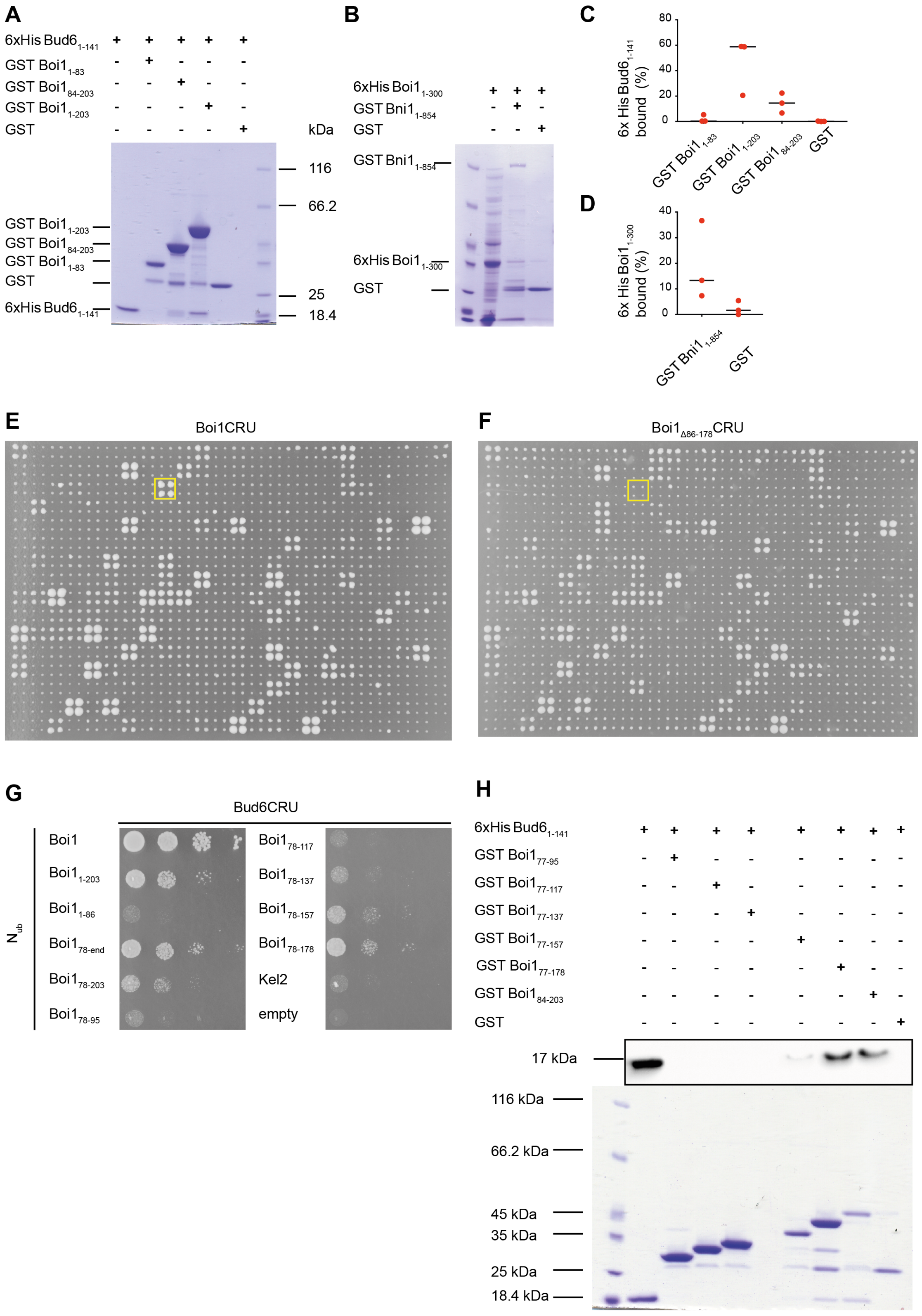


Defining the interaction sites between Boi1 and Bud6 and Bni1. (A-B) Coomassie-staining of SDS-PAGEs corresponding to western blots shown in Fig.1F. (C-D) Quantitative analysis of three independent pull down experiments of the representative examples shown in Fig.1F. The fraction of bound proteins were determined relative to the loaded input and based on signal intensities from anti-His western blots. See Materials and Methods section for experimental details. (E, F) as in Fig. S1, showing the complete arrays corresponding to the cut-outs shown in Fig. 1G. Cells were incubated for 3 days on medium containing 5-FOA and 100 µM copper. (G) Haploid yeast cells containing Bud6CRU and co-expressing the indicated $N_{ub}$ fusion protein were spotted in serial dilutions starting with an $OD_{600}$=1 on medium containing FOA and 100 µM copper, and incubated for 2 days. $N_{ub}$-Kel1 should not interact with Bud6CRU. (H) Pull down analysis. Purified 6xHisBud6$_{1-141}$ was tested against GST, or different GST-fragments of Boi1. Upper panel shows the anti-His western blot after SDS-PAGE and transfer on nitrocellulose of the glutathione-eluates of the GST-fusion-displaying beads. Lower panel shows the coomassie-staining of the same samples but loaded on a different gel. Assay was performed as a triplicate with similar outcome.

#### Figure S3

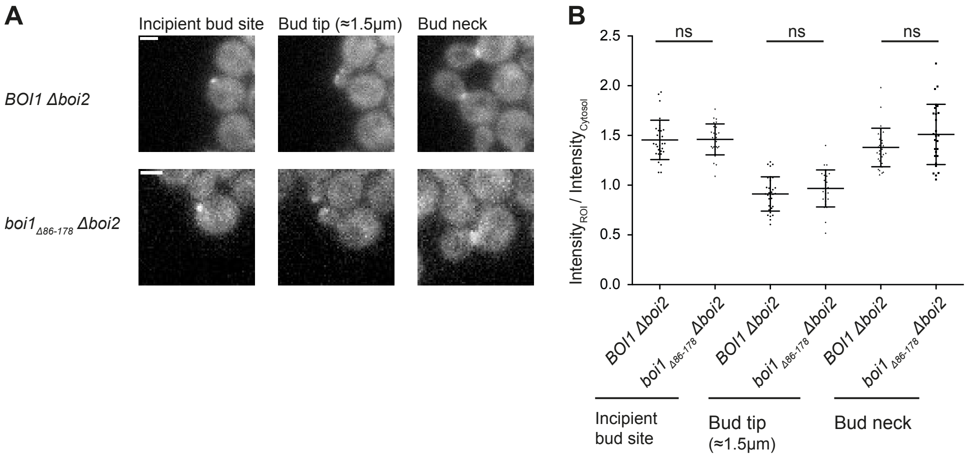


Bud6 localizes to sites of polar growth independently of Boi1/2. (A) Stills of a time-lapse microscopy of *BOI1 Δboi2-,* and *boi1*$_{Δ86-178}$ *Δboi2*-cells expressing Bud6-GFP. Representative images show maximum projections of 5 stacks. Scale bar indicates 2 µm. (B) Quantification of the relative fluorescence enrichment of Bud6-GFP at the incipient bud site, bud tip, and bud neck, normalized to the cytosolic mean intensity of the mother cell after background-subtraction. Fluorescence microscopy was performed in two independent measurements including in total $n_{Δboi2}$= 31, $n_{boi1Δ86-178Δboi2}$= 27 cells per time point. Statistical significance was determined by a Kruskal-Wallis test, followed by Dunn´s multiple comparison. ns= not significant.

### Figure S4

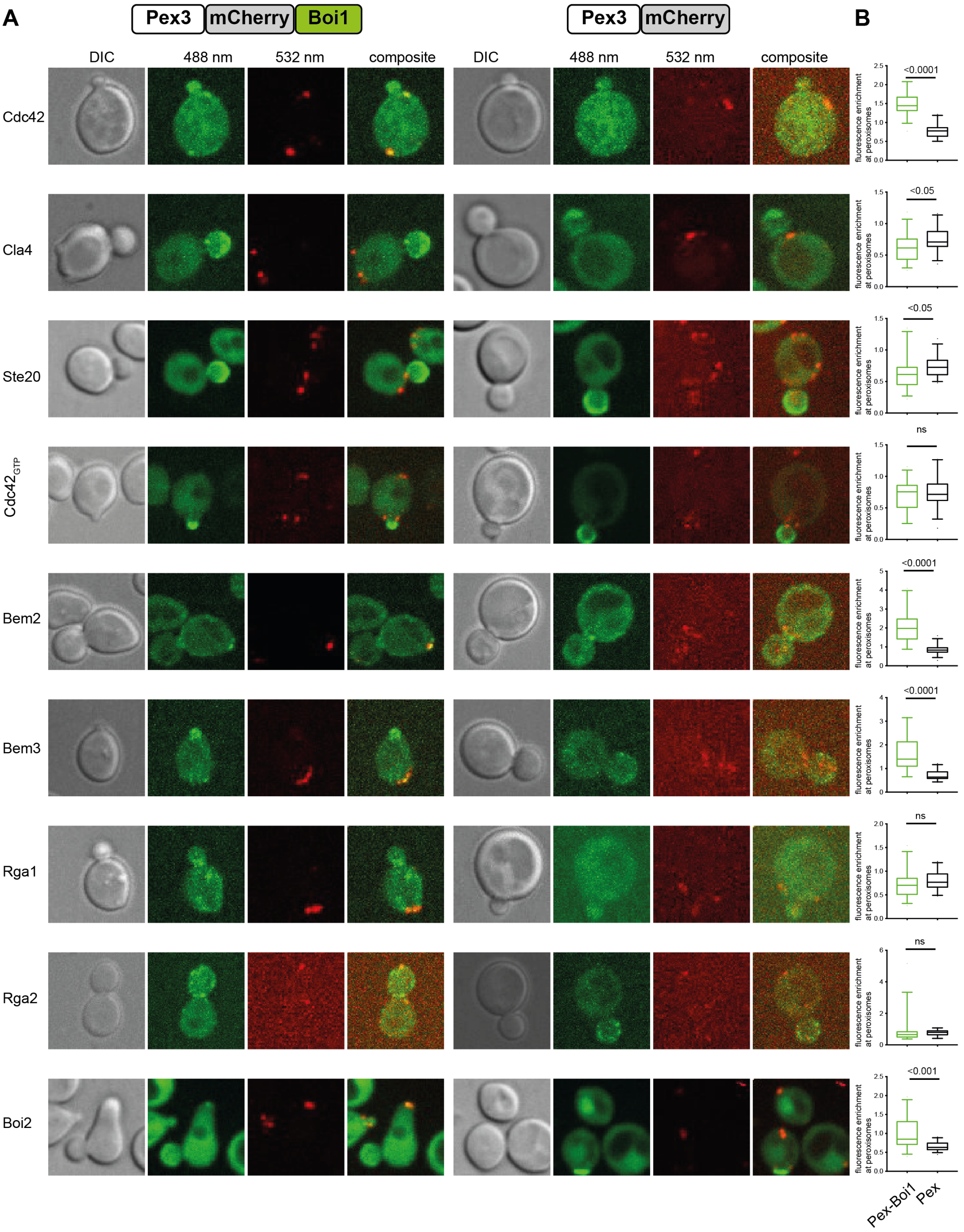
A
Pex3
mCherry
Boi1
Pex3
mCherry
B
DIC
488 nm
532 nm
composite
DIC
488 nm
532 nm
composite
Cdc42
Cla4
Ste20
Cdc42GTP
Bem2
Bem3
Rga1
Rga2
Boi2
<0.0001
<0.05
<0.05
ns
<0.0001
<0.0001
ns
ns
<0.001
fluorescence enrichment at peroxisomes
Pex-Boi1
Pex

#### Figure S4 continued

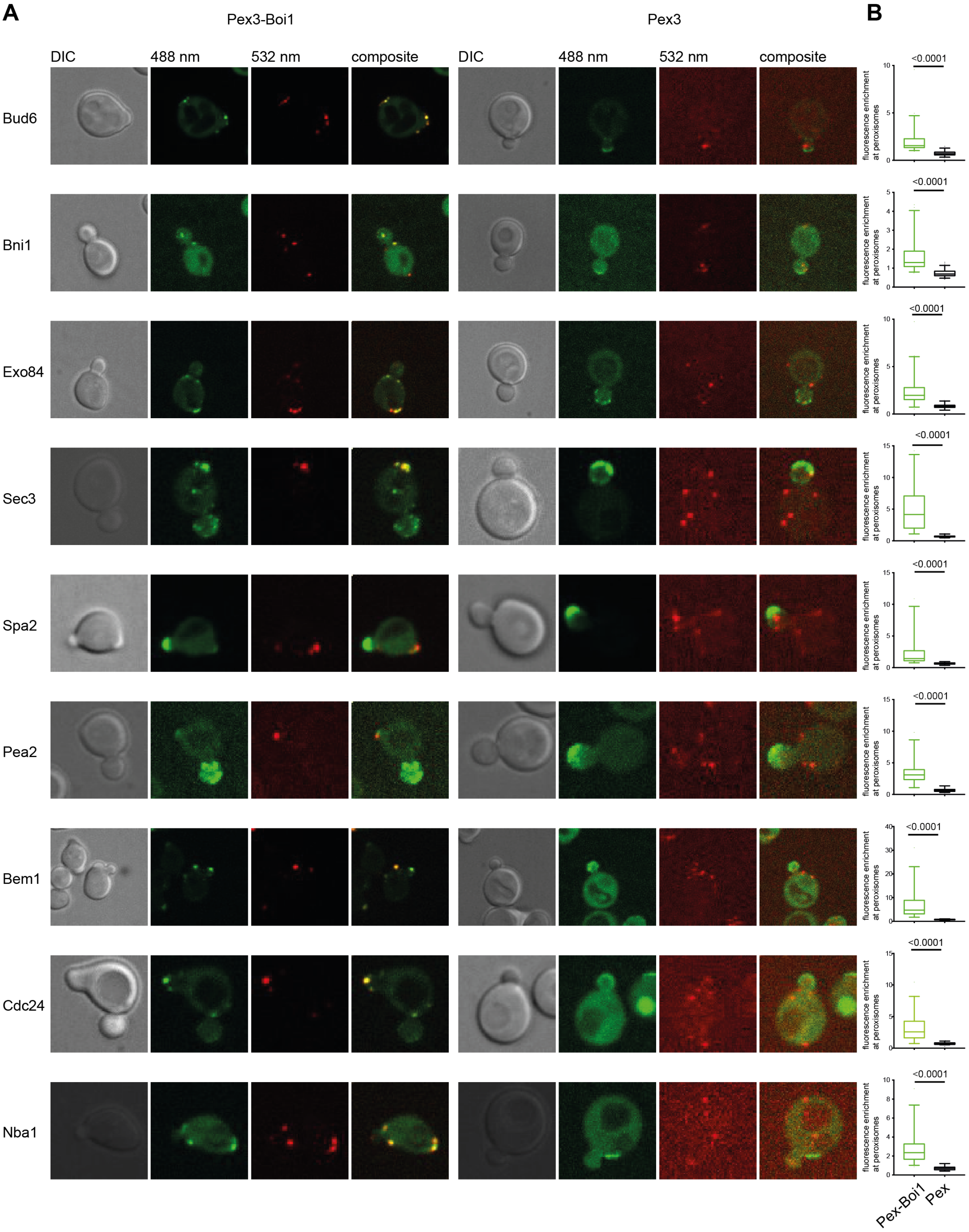


Recruitment of polarity proteins to Boi1-decorated peroxisomes. (A) Images of cells co-expressing Pex3$_{1\text{-}45}$mCherry, or Pex3$_{1\text{-}45}$mCherryBoi1 together with the indicated GFP fusion proteins. Images show the maximum projections of 10 stacks. The images of cells expressing Pex3-Boi1 together with Bud6-, Bni1-, Exo84-, and Sec3-GFP are taken from Fig. 4C. Active Cdc42 (Cdc42$_{GTP}$) was probed with Gic2-CRIB-GFP. (B) The relative fluorescence enrichments of the GFP fusion proteins at Pex3$_{1\text{-}45}$mCherry-, or Pex3$_{1\text{-}45}$mCherryBoi1-decorated peroxisomes relative to the cytosol were quantified in at least 30 cells deriving from two independent measurements. Boxplot with 5 to 95 % whiskers are shown. Man-Whitney tests were used for evaluating the statistical significances of the measured differences. Ns= not significant.

**Supplementary Tables**

**Supplementary Table S1**

Split-Ub interaction partners of Bni1 and Bud6.

| $N_{ub}$ \ CRU | Bud6 | Bni1 |
|---|---|---|
| Aip1 | + | - |
| Bil1 | + | + |
| Bni5 | + | - |
| Boi1 | + | + |
| Boi2 | + | + |
| Bud4 | + | - |
| Bud6 | + | + |
| Bud14 | + | - |
| Chc1 | + | + |
| Chs1 | + | - |
| Cnb1 | - | + |
| Crm1 | - | + |
| Ede1 | + | + |
| Epo1 | + | - |
| Exo70 | - | + |
| Mlc1 | - | + |
| Msb1 | + | - |
| Nba1 | + | - |
| Nis1 | + | - |
| Nsp1 | + | - |
| Osh6 | - | + |
| Pea2 | + | - |
| Sac6 | + | + |
| Sec4 | + | - |
| Sec5 | - | + |
| Sec8 | - | + |
| She2 | - | + |
| Spa2 | + | - |
| Vma13 | + | - |
| YFR016c/Aip5 | + | - |

Interactions were obtained by analyzing the large-scale Split-Ub assays shown in Figure S1. (+) or (-) indicate the presence or absence of an interaction.

**Supplementary Table S2**

Quantification of the local enrichments of GFP-labeled proteins at Pex-Boi1-, Pex-Boi1$_{\Delta 86-178}$-, or Pex-decorated peroxisomes.

| **Protein** | $I_{Pex\text{-}Boi1}$ | $I_{Pex}$ | $I_{Pex\text{-}Boi1\Delta 86\text{-}178}$ | $I_{Pex\text{-}Boi1}/I_{Pex}$ |
|---|---|---|---|---|
| Bem1 | 7.00 (6.20) | 0.73 (0.16) | 1.55 (1.17) | 9.54 |
| Bem2 | 2.02 (0.78) | 0.85 (0.24) | n.d. | 2.37 |
| Bem3 | 1.67 (0.75) | 0.72 (0.21) | n.d. | 2.33 |
| Bni1 | 1.68 (0.91) | 0.73 (0.20) | 1.23 (0.32) | 2.29 |
| Boi2 | 0.97 (0.40) | 0.66 (0.11) | n.d. | 1.47 |
| Bud6 | 1.98 (1.37) | 0.72 (0.24) | 0.67 (0.25) | 2.76 |
| Cdc24 | 3.24 (2.23) | 0.74 (0.19) | 2.44 (1.97) | 4.39 |
| Cdc42 | 1.47 (0.28) | 0.78 (0.18) | n.d. | 1.89 |
| Cla4 | 0.61 (0.21) | 0.76 (0.20) | n.d. | 0.81 |
| Exo84 | 2.37 (1.63) | 0.82 (0.24) | 1.72 (0.95) | 2.90 |
| Cdc42$_{GTP}$ | 0.71 (0.22) | 0.77 (0.25) | n.d. | 0.92 |
| Nba1 | 2.74 (1.67) | 0.72 (0.22) | n.d. | 3.78 |
| Pea2 | 3.55 (1.97) | 0.70 (0.28) | n.d. | 5.07 |
| Rga1 | 0.72 (0.27) | 0.80 (0.19) | n.d. | 0.89 |
| Rga2 | 0.89 (0.90) | 0.77 (0.18) | n.d. | 1.15 |
| Sec3 | 5.29 (4.06) | 0.69 (0.15) | 1.62 (0.68) | 7.66 |
| Spa2 | 2.51 (2.55) | 0.65 (0.16) | n.d. | 3.86 |
| Ste20 | 0.64 (0.24) | 0.88 (0.44) | n.d. | 0.72 |

Relative fluorescence enrichments of GFP-labeled proteins at peroxisomes in *Δboi1*-cells expressing $P_{MET17}$Pex3$_{1-45}$mCherry/-Boi1/-Boi1$_{\Delta 86-178}$. The ratios of the fluorescence intensities of Pex3-Boi1- to Pex3-decorated peroxisomes are shown in the right lane. Values in brackets indicate the s.d.. Active Cdc42 (Cdc42$_{GTP}$) was probed with Gic2-CRIB-GFP. n.d. = not determined

**Supplementary Table S3**

List of primers used in this study

| Primer name | Sequence 5´->3 |
|---|---|
| Bud6 Eag CUB | CCTCCCCGGCCGCGTATTCTCCTCGAGCAAATTC |
| Bud6 Sal CUB | CCACCCGTCGACCCAGTAAACCCCGGCCCAAAATA |
| Bud6 CUB Ctr | CGCATTAAGGCTCCTATTTAT |
| Bni1_EAG_CUB | CCTCCCGGCCGAACGCTCCACAATTAC |
| Bni1_SAL_CUB | CCTCCGTCGACCCTTTGAAACTTAGCCTGTTA |
| Bni1_Cub CTR new | CCAGAAAGAGAGCCCTAGTGAGG |
| Boi1 Eag Cub | CCTCCCGGCCGAAGAACAGTAGCTCCAAAGATGG |
| Boi1 Sal Cub | CCACCGTCGACCCAAGTCTGCCACCAGGGTATTC |
| Boi1 Cub CTR 870 | GCACAAGCACAAGAGCAAGCAC |
| Cub ctr | TCTTCTAGCTGCTTACCG |
| GFP (S65T) ATG+84RV | CGC CTT CAC CCT CTC CAC TGA C |
| pGAL1 prom RV | CGGAGGGCTGTCACCCGCTC |
| KlURA3prom RV | GCGTTACCACCATCCAATGC |
| Boi1 aa1 Eag Fw | CCTCCCGGCCGATGAGTCTCGAAGGAAATACC |
| Boi1 155 RV EcoRI | GACCGGAATTCCCTTCGGGCACGCGTGTGG |
| Boi1203NuiAcc65I | CCTCCGGTACCTGACTTACTTATATTCTTCAAAG |
| Boi1 Nub fw | CTAACTCGAGGTGACCGGAGGCCACTGTAATAATAAAAAATAGAAGGCATAGGCCACTAGTGGATC |
| Nub Boi1D203 RV | GACGAACTGATTCCGAACTCGGTTCTAGGGATTTTGTTGATATATTGGTCGACCCCGCATAGTCAGG |
| Nub Boi1D300 RV | GTTGGGCAGCTTGGTTTTCAAAAGCAAAGGTAGAGTAGTCGGCGTCGGTCGACCCCGCATAGTCAGG |
| Nub Boi1D416 RV | GTTGTTTGAGGAGGAAACCTTGCGTTATTATTTGTGTACCTGTTATTGGTCGACCCCGCATAGTCAGG |
| gR1 Boi1-304 | GATCAATGATTCTGCGAGTAACATGTTTTAGAGCTAG |
| gR2 Boi1-304 | CTAGCTCTAAAACATGTTACTCGCAGAATCATT |
| DepeI Boi1 131 AS | CTCTTGTTCAACTGAACCACTTCTTAGCTCTTCCAAGGCTTTGTCTTCGTACGCTGCAGGTCGAC |
| DepeII Boi1 131 AS | CAGGAGAACAGATATACATCATTGAAAAGTACAATGAGCGATATATAGGGATAACAGGGTAATCCGCGCGTTGGCCGATTCAT |
| DepeIII Boi1 86-178 | ATAGAAAAACCAGAGAACCTGCACAAATCAACACATGACTCGTTATTTTCTAGCACAGCG |
| S1 Boi1 | AGTTCTAACTCGAGGTGACCGGAGGCCACTGTAATAATAAAAAATAGAAGCGTACGCTGCAGGTCGAC |
| S2 Boi1 | ATTAAGGTGTTTAAGTTGGTCAAGAAGTAACTAATGATTGCAGTTC ATC GAT GAA TTC GAG CTC G |
| G1 Boi1 | TTG ACG CGT AAG AAA GGC TAT |
| G2 Boi1 | CAG CCC ATT ATT TGC TGG GTC |
| S1 Boi2 | TTATTGAATCATACCAACTTCTCCGCAAAAAATTTGAACATATAAAGCGTACGCTGCAGGTCGAC |
| S2 Boi2 | TAGCTTAATGAGTTTATGCATCAAATTTGAGGCGC |

| Primer name | Sequence 5´->3 |
|---|---|
| | ATCTTTTTCAATAGCTTTAAAAATCGATGAATTCGAGCTCG |
| G1 Boi2 | CTATTTGCGTAATTCCCTTCTGC |
| G2 Boi2 | AAGGATCATCGGCATCCTTTC |
| H3 | GGAAGGAGTTAGACAACCTG |
| AgTefpRV | CATGTCGCTGGCCGGGTGAC |
| Boi1_Glu78_SalI_fw | CGAT GTCGAC C GAAAAACCAGAGAACCTGC |
| Boi1_Gly95_Stop_Acc65I_rv | CGAT GGTACC TTA ACCAGAATTTCCACTCTCTTG |
| Boi1_Glu117_Stop_Acc65I_rv | CGAT GGTACC TTA CTCCTGTTGATGCGAGGAG |
| Boi1_Leu137_Stop_Acc65I_rv | CGAT GGTACC TTA TAGCTCTTCCAAGGCTTTG |
| Boi1_Ser157_Stop_Acc65I_rv | CGAT GGTACC TTA GCTAACTTCGGGCACGC |
| Boi1_Thr178_Stop_Acc651_rv | CGAT GGTACC TTA CGTGTTTTCCTCATTTCTGG |
| Boi1_Glu77_NcoI_fw | CCTCCCCATGGCCGAAAAACCAGAGAACCTGCAC |
| Boi1_Gly95stop_EcoRI_rv | CCTCCGAATTCTTAACCAGAATTTCCACTCTCTTG |
| Boi1_Glu117_Stop_EcoRI_rv | CCTCCGAATTCTTACTCCTGTTGATGCGAGGAG |
| Boi1_Leu137stop_EcoRI_rv | CCTCCGAATTCTTATAGCTCTTCCAAGGCTTTGTC |
| Boi1_Ser157stop_EcoRI_rv | CCTCCGAATTCTTAGCTAACTTCGGGCACGCG |
| Boi1_Thr178stop_EcoRI_rv | CCTCCGAATTCTTACGTGTTTTCCTCATTTCTGG |

**Supplementary Table S4**

Yeast and *E.Coli* strains used in this study.

| Strain | Description | Source | Identifier |
|---|---|---|---|
| *E.coli* XL1 blue | *endA1 gyrA96(NalR) thi-1 recA1 relA1 lac glnV44 F' [::Tn10 proAB+ lacIq (lacZ)M15] hsdR17(rK- mK+)* | Stratagene | XL1 blue |
| *E.coli* Bl21 DE3 | *F- ompT gal dcm lon hsdSB( rB-mB-) λ(DE3) pLysS(CmR)* | Invitrogen | Bl21DE3 |
| Yeast JD47 | *MATa, his3-Δ200, leu2-3, 112 lys2-801, trp1-Δ63, ura3-52* | (Dohmen et al. 1995) | JD47 |
| Yeast JD53 | *MATα, his3-Δ200, leu2-3, 112 lys2-801, trp1-Δ63, ura3-52* | (Dohmen et al. 1995) | JD53 |
| JD47 BUD6CRU | *BUD6::BUD6-CRU, HIS3* | This study | YWY43 |
| JD47 BNI1CRU | *BNI1::BNI1-CRU, HIS3* | This study | SY229 |
| JD47 BOI1CRU | *BOI1::BOI1-CRU, HIS3* | (Kustermann et al. 2017) | NJY317 |
| JD47 $BOI1_{Δ86\text{-}178}$ CRU | $BOI1_{Δ86\text{-}178}$*::*$BOI1_{Δ86\text{-}178}$ *–CRU, HIS3* | This study | YOG121 |
| JD53 $N_{ub}$-Boi1 *ΔBOI2* | $P_{BOI1}$*::kanMX6* $P_{CUP1}N_{ub}$*-HA, BOI2::hph* | This study | YOG178 |
| JD53 $N_{ub}$-Boi1 $_{Δ203,}$ *ΔBOI2* | $P_{BOI1}BOI1_{1\text{-}203}$*::kanMX6* $P_{CUP1}N_{ub}$*-HA, BOI2::hph* | This study | YOG180 |
| JD53 $N_{ub}$-Boi1 $_{Δ299,}$ *ΔBOI2* | $P_{BOI1}BOI1_{1\text{-}299}$*::kanMX6* $P_{CUP1}N_{ub}$*-HA, BOI2::hph* | This study | YOG182 |
| JD53 $N_{ub}$-Boi1 $_{Δ414,}$ *ΔBOI2* | $P_{BOI1}BOI1_{1\text{-}414}$*::kanMX6* $P_{CUP1}N_{ub}$*-HA, BOI2::hph* | This study | YOG183 |
| JD47 *ΔBOI1ΔBOI2* $P_{MET17}$SSO1 mCherry | *BOI1::hph, BOI2::nat,* $P_{MET17}$*SSO1-mCherry pRS315 LEU2* | This study | YOG203 |
| JD47 $P_{Gal1}$ BOI1-mCherry, BUD6-GFP | $P_{BOI1}$*::kanMX6* $P_{GAL1}$*BOI1::BOI1-mCherry URA3, BUD6::BUD6 GFP TRP1* | This study | YWY302 |
| JD47 $P_{Gal1}$ BOI1-mCherry, BNI1-3GFP | $P_{BOI1}$*::kanMX6* $P_{GAL1}$*BOI1::BOI1-mCherry URA3, BNI1::BNI1 3GFP TRP1* | This study | YWY303 |
| JD47 $P_{Gal1}$BOI1-mCherry, Lifeact GFP | $P_{BOI1}$*::kanMX6* $P_{GAL1}$*BOI1::BOI1-mCherry URA3, CEN* $P_{ABP140}ABP140_{1\text{-}17}$ *HIS3* | This study | YWY301 |
| JD53 $N_{ub}$-Boi1 *ΔBOI2*, GFP-Sec4 | $P_{BOI1}$*::kanMX6* $P_{CUP1}N_{ub}$*-HA, BOI2::hph, ura3-52::URA3* $P_{SEC4}$*GFP-SEC4Term*$_{SEC4}$ | This study | YOG226 |

| Strain | Description | Source | Identifier |
|---|---|---|---|
| JD53 $N_{ub}$-Boi1 $_{\Delta 203}$, *ΔBOI2*, GFP-Sec4 | *BOI1$_{1-203}$::kanMX6 $P_{CUP1}N_{ub}$-HA BOI2::hph, ura3-52::URA3 $P_{SEC4}$GFP-SEC4Term$_{SEC4}$* | This study | YOG228 |
| JD53 $N_{ub}$-Boi1 $_{\Delta 299}$, *ΔBOI2*, GFP-Sec4 | *BOI1$_{1-299}$::kanMX6 $P_{CUP1}N_{ub}$-HA BOI2::hph, ura3-52::URA3 $P_{SEC4}$GFP-SEC4Term$_{SEC4}$* | This study | YOG230 |
| JD53 $N_{ub}$-Boi1 $_{\Delta 414}$, *ΔBOI2*, GFP-Sec4 | *BOI1$_{1-414}$::kanMX6 $P_{CUP1}N_{ub}$-HA BOI2::hph, ura3-52::URA3 $P_{SEC4}$GFP-SEC4Term$_{SEC4}$* | This study | YOG232 |
| JD53 Nub Boi1$_{\Delta 86-178}$ | *BOI1:: kanMX6 $P_{CUP1}N_{ub}$-HA BOI1$_{\Delta 86-178}$* | This study | YOG535 |
| JD47 *ΔBOI2* | *BOI2::hph* | This study | YOG162 |
| JD47 *ΔBOI2* BOI1$_{\Delta 86-178}$ | *BOI2::hph, BOI1::BOI1$_{\Delta 86-178}$* | This study | YOG126 |
| JD47 *ΔBOI1*, $P_{MET17}$Pex3$_{1-45}$mCherryBoi1 | *BOI1::nat, ura3-52::URA3 $P_{MET17}$PEX3$_{1-45}$mCherryBOI1* | This study | YOG239 |
| JD47 *ΔBOI1*, $P_{MET17}$Pex3$_{1}$-45-mCherry | *BOI1::nat, ura3-52::URA3 $P_{MET17}$PEX3$_{1-45}$mCherry* | This study | YOG237 |
| JD47 *ΔBOI1*, $P_{MET17}$Pex3$_{1-45}$mCherryBoi1$_{\Delta 86-178}$ | *BOI1::nat, ura3-52::URA3 $P_{MET17}$PEX3$_{1-45}$mCherryBOI1$_{\Delta 86-178}$* | This study | YOG245 |
| JD47 *ΔBOI1*, $P_{MET17}$Pex3$_{1-45}$mCherryBoi1, $P_{MET17}$GFP-Sec4 (cen) | *BOI1::nat, ura3-52::URA3 $P_{MET17}$PEX3$_{1-45}$mCherryBOI1, CEN $P_{MET17}$GFP-SEC4 HIS3* | This study | YOG251 |
| JD47 *ΔBOI1*, $P_{MET17}$Pex3$_{1-45}$mCherry $P_{MET17}$GFP-Sec4 (cen) | *BOI1::nat, ura3-52::URA3 $P_{MET17}$PEX3$_{1-45}$mCherry CEN $P_{MET17}$GFP-SEC4 HIS3* | This study | YOG256 |
| JD47 *ΔBOI1*, $P_{MET17}$Pex3$_{1-45}$mCherryBoi1$_{\Delta 86-178}$ $P_{MET17}$GFP-Sec4 (cen) | *BOI1::nat, ura3-52::URA3, $P_{MET17}$PEX3$_{1-45}$mCherryBOI1$_{\Delta 86-178}$, CEN $P_{MET17}$GFP-SEC4 HIS3* | This study | YOG289 |
| JD47 $P_{MET17}$GFPSec4 | *ura3-52::URA3, $P_{MET17}$GFP-SEC4* | This study | YOG540 |
| JD47 *ΔBOI2* $P_{MET17}$GFP-Sec4 | *BOI2::hph, ura3-52::URA3, $P_{MET17}$ GFP-SEC4* | This study | YOG541 |
| JD47 *ΔBOI2* BOI1 $_{\Delta 86-178}$ $P_{MET17}$GFP-Sec4 | *BOI1::BOI1 $_{\Delta 86-178}$, BOI2::hph, ura3-52::URA3, $P_{MET17}$GFP-SEC4* | This study | YOG542 |

**Supplementary Table S5**

List of plasmids used in this study.

| Plasmid | Description | Source |
|---|---|---|
| pAC | $P_{T7}$*6HIS, kanMX*$^R$ | (Iffland et al. 2000) |
| 6xHis Bud6$_{1\text{-}141}$ | $P_{T7}$*6HIS-BUD6*$_{1\text{-}141}$*, kanMX*$^R$ | This study |
| 6xHis Boi1$_{1\text{-}300}$ | $P_{T7}$*6HIS-BOI1*$_{1\text{-}300}$*, kanMX*$^R$ | This study |
| 6xHis Bud6$_{1\text{-}364}$ | $P_{T7}$*6HIS-BUD6*$_{1\text{-}364}$*, kanMX*$^R$ | This study |
| MBP Bud6$_{550\text{-}688}$ | $P_{tac}$*MBP-BUD6*$_{550\text{-}688}$*, Amp*$^R$ | This study |
| pMal c5x | $P_{tac}$*MBP, Amp*$^R$ | New England Biolabs |
| pGex2T | $P_{tac}$*GST, Amp*$^R$ | GE Healthcare |
| GST Boi1$_{1\text{-}83}$ | $P_{tac}$*GST-BOI1*$_{1\text{-}83}$*,Amp*$^R$ | This study |
| GST Boi1$_{84\text{-}203}$ | $P_{tac}$*GST-BOI1*$_{84\text{-}203}$*,Amp*$^R$ | This study |
| GST Boi1$_{1\text{-}203}$ | $P_{tac}$*GST-BOI1*$_{1\text{-}203}$*,Amp*$^R$ | This study |
| GST Bni1$_{1\text{-}854}$ | $P_{tac}$*GST-BNI1*$_{1\text{-}854}$*,Amp*$^R$ | This study |
| GST Boi1$_{77\text{-}95}$ | $P_{tac}$*GST-BOI1*$_{77\text{-}95}$*,Amp*$^R$ | This study |
| GST Boi1$_{77\text{-}117}$ | $P_{tac}$*GST-BOI1*$_{77\text{-}117}$*,Amp*$^R$ | This study |
| GST Boi1$_{77\text{-}137}$ | $P_{tac}$*GST-BOI1*$_{77\text{-}137}$*,Amp*$^R$ | This study |
| GST Boi1$_{77\text{-}157}$ | $P_{tac}$*GST-BOI1*$_{77\text{-}157}$*,Amp*$^R$ | This study |
| GST Boi1$_{77\text{-}178}$ | $P_{tac}$*GST-BOI1*$_{77\text{-}178}$*,Amp*$^R$ | This study |
| Cub-R-Ura3-pRS 303 empty | *pRS 303, lacZ, Amp*$^R$*, His3* | (Wittke et al. 1999) |
| Bud6CRU pRS303 | *Bud6-Cub-R-Ura3 pRS303* | This study |
| Bni1CRU pRS303 | *Bni1-Cub-R-Ura3 pRS303* | This study |
| Boi1CRU pRS303 | *Boi1-Cub-R-Ura3 pRS303* | (Kustermann et al. 2017) |
| Bud6-GFP, pRS304 (Linearization with EcoRI) | *Bud6-GFP, pRS304, TRP1, Amp*$^R$ | (Glomb, Bareis, Johnsson 2019) |
| Bni1-GFP, pRS304 (Linearization with ApaI) | *Bni1-GFP, pRS304, TRP1, Amp*$^R$ | This study |
| Bem1-GFP, pRS304 (Linearization with BsmI) | *Bem1-GFP, pRS304, TRP1, Amp*$^R$ | This study |
| Exo84-GFP, pRS304 (Linearization with Bgl II) | *Exo84-GFP, pRS304, TRP1, Amp*$^R$ | This study |
| Nba1-GFP, pRS304 (Linearization with SpeI) | *Nba1-GFP, pRS304, TRP1, Amp*$^R$ | This study |
| Pea2-GFP, pRS304 (Linearization with EcoRI) | *Pea2-GFP, pRS304, TRP1, Amp*$^R$ | This study |
| Spa2-GFP, pRS304 (Linearization with *ClaI*) | *Spa2-GFP, pRS304, TRP1, Amp*$^R$ | This study |
| Cdc24-GFP, pRS304 (Linearization with Bgl II) | *CDC24-GFP, pRS304, TRP1, Amp*$^R$ | This study |

| Plasmid | Description | Source |
|---|---|---|
| Sec3-GFP, pRS304 (Linearization with Eco91I) | *SEC3-GFP, pRS304, TRP1, $Amp^R$* | This study |
| Cla4-GFP, pRS304 (Linearization with XhoI) | *CLA4-GFP, pRS304, TRP1, $Amp^R$* | This study |
| Ste20-GFP, pRS304 (Linearization with Eco47III) | *STE20-GFP, pRS304, TRP1, $Amp^R$* | This study |
| $P_{GIC2}GIC2_{1-208}$-CRIB ($Cdc42_{GTP}$) GFP, pRS314 | *$P_{GIC2}GIC2_{1-208}$ CRIB-GFP, CEN, pRS314, TRP1, $Amp^R$* | This study |
| Bem2-GFP, pRS304 (Linearization with BsmI) | *BEM2-GFP, pRS304, TRP1, $Amp^R$* | This study |
| Bem3-GFP, pRS304 (Linearization with BsaBI) | *BEM3-GFP, pRS304, TRP1, $Amp^R$* | This study |
| Rga1-GFP, pRS304 (Linearization with EcoRI) | *RGA1-GFP, pRS304, TRP1, $Amp^R$* | This study |
| Rga2-GFP, pRS304 (Linearization with EcoRI) | *RGA2-GFP, pRS304, TRP1, $Amp^R$* | This study |
| Boi2-GFP, pRS304 (Linearization with SpeI) | *BOI2-GFP, pRS304, TRP1, $Amp^R$* | This study |
| GFP Cdc42, pRS314 | *$P_{CDC42}$GFP-CDC42, TRP1, pRS314, $Amp^R$* | This study |
| Boi1-mCherry, pRS306 (Linearization with AgeI) | *BOI1-mCherry, pRS306, URA3* | This study |
| $P_{MET17}$-$Pex3_{1-45}$-mCherry-mcs, pRS306 | *$P_{MET17}$-$PEX3_{1-45}$-mCherry-mcs, pRS306, URA3, $Amp^R$* | (Glomb, Bareis, Johnsson 2019) |
| $P_{MET17}$-$Pex3_{1-45}$-mCherry-Boi1, pRS306 | *$P_{MET17}$-$PEX3_{1-45}$-mCherry-BOI1, pRS306, URA3, $Amp^R$* | This study |
| $P_{MET17}$-$Pex3_{1-45}$-mCherry-$Boi1_{\Delta 86-178}$, pRS306 | *$P_{MET17}$-$PEX3_{1-45}$-mCherry-$BOI1_{\Delta 86-178}$, pRS306, URA3, $Amp^R$* | This study |
| pGSKU | *$Amp^R$, KanMX4, KlURA3, GAL1-I-SceI* | (Storici and Resnick 2006) |
| pFA hphNT1 | *pAgTEF-hphNT1, $Amp^R$* | (Janke et al. 2004) |
| pFA natNT2 | *pAgTEF-natNT2, $Amp^R$* | (Janke et al. 2004) |
| pFA CmLEU2 | *pAgTEF-CmLEU2, $Amp^R$* | (Bahler et al. 1998) |
| pML104 | *backbone: pRSII426* | (Laughery et al. 2015) |
| pML104-BOI1-304 | *pML104* | This study |
| $P_{CUP1}$NuiHA-kanMX4 | *$P_{CUP1}$NuiHA-kanMX4, $Amp^R$* | (Hruby et al. 2011) |
| $P_{SEC4}$GFP-Sec4 pRS306 (Linearization with ApaI) | *$P_{Sec4}$ GFP-Sec4 $Term_{Sec4}$ pRS306, URA3, $Amp^R$* | This study, based on (Donovan and Bretscher 2015) |

| Plasmid | Description | Source |
|---|---|---|
| $P_{MET17}$GFP-Sec4 pRS306 (Linearization with StuI) | *$P_{MET17}$ GFP-SEC4 $Term_{Sec4}$ pRS306, URA3, $Amp^R$* | This study |
| $P_{MET17}$GFP-Sec4, pRS313 | *$P_{MET17}$GFP-SEC4, HIS3, $Amp^R$, pRS313* | This study |
| Lifeact GFP, pRS313 | *$P_{ABP140}$ABP140$_{1-17}$-GFP, HIS3* | This study, based on (Riedl et al. 2008) |
| $N_{ub}$-Guk1 | *$P_{CUP1}N_{ub}$-HA-GUK1 kanMX4 CEN, $Amp^R$* | (Kustermann et al. 2017) |
| $N_{ub}$- (empty) | *$P_{CUP1}N_{ub}$-HA kanMX4 CEN, $Amp^R$* | (Hruby et al. 2011) |
| $N_{ub}$-Bud6$_{1-364}$ cen | *$P_{CUP1}N_{ub}$-HA kanMX BUD6$_{1-364}$, CEN* | (Glomb, Bareis, Johnsson 2019) |
| $N_{ub}$-Bud6 $_{360\text{-end}}$ cen | *$P_{CUP1}N_{ub}$ HA kanMX BUD6$_{360-788}$, CEN* | (Glomb, Bareis, Johnsson 2019) |
| $N_{ub}$-Bud6$_{1-240}$ cen | *$P_{CUP1}N_{ub}$-HA kanMX BUD6$_{1-240}$, CEN* | (Glomb, Bareis, Johnsson 2019) |
| $N_{ub}$-Bud6$_{1-141}$ cen | *$P_{CUP1}N_{ub}$ HA kanMX BUD6$_{1-141}$, CEN* | (Glomb, Bareis, Johnsson 2019) |
| $N_{ub}$-Boi1$_{78-95}$ cen | *$P_{CUP1}N_{ub}$ HA kanMX BOI1$_{78-95}$, CEN* | This study |
| $N_{ub}$-Boi1$_{78-117}$ cen | *$P_{CUP1}N_{ub}$ HA kanMX BOI1$_{78-117}$, CEN* | This study |
| $N_{ub}$-Boi1$_{78-137}$ cen | *$P_{CUP1}N_{ub}$ HA kanMX BOi1$_{78-137}$, CEN* | This study |
| $N_{ub}$-Boi1$_{78-157}$ cen | *$P_{CUP1}N_{ub}$-HA kanMX BOI1$_{78-157}$, CEN* | This study |
| $N_{ub}$-Boi1$_{78-178}$ cen | *$P_{CUP1}N_{ub}$-HA kanMX BOI$_{78-178}$, CEN* | This study |
| $P_{MET17}$Sso1-mCherry, pRS315 | *$P_{MET17}$ SSO1-mCherry, CEN, LEU2, $Amp^R$* | This study |
| $P_{MET17}$GFP-Snc1 pRS313 | *$P_{MET17}$GFP-SNC1, CEN, HIS3, $Amp^R$* | This study |

**Supplementary Table S6**

Conditions for protein expression.

| Protein | Expression condition (IPTG concentration, expression temperature, -duration, -medium) |
|---|---|
| 6xHis-Bud6$_{1-141}$ | 1 mM IPTG, 37°C, 5h, LB |
| 6xHis-Boi1$_{1-300}$ | 0.1 mM IPTG, 37°C, 6h, LB |
| GST | 1 mM IPTG, 37 °C, 4h, LB |
| GST-Boi1$_{1-83}$ | 0.1 mM IPTG, 18°C, o.n., SB |
| GST-Boi1$_{1-203}$ | 1 mM IPTG, 37°C, 4h, LB |
| GST-Boi1$_{84-203}$ | 0.1 mM IPTG, 18°C, 6h, SB |
| GST-Bni1$_{1-854}$ | 1 mM IPTG, 18°C, o.n., SB |
| GST-Boi1$_{77-95}$ | 0.1 mM IPTG, 18°C, 6h, SB |
| GST-Boi1$_{77-117}$ | 0.1 mM IPTG, 18°C, 6h, SB |
| GST-Boi1$_{77-137}$ | 0.1 mM IPTG, 18°C, 6h, SB |
| GST-Boi1$_{77-157}$ | 0.1 mM IPTG, 18°C, 6h, SB |
| GST-Boi1$_{77-178}$ | 0.1 mM IPTG, 18°C, 6h, SB |

#### Supplementary videos

##### Video S1

Impairing the Bud6-Boi1 interaction disturbs directed secretory vesicle movement in the bud. GFP-Sec4 labeled secretory vesicles were followed in cells of the indicated genotypes over a time course of 103 s, taking an image of 5 z stacks every second, and stacked to maximum projections of the middle 3 stacks.

##### Video S2

Boi1-decorated peroxisomes attract secretory vesicles.
Secretory vesicles were tracked over a time course of 35 s, taking an image every 100 ms of a single z layer in *Δboi1* cells expressing Pex3/-Boi1/-Boi1$_{\Delta 86\text{-}178}$. Before starting time-lapse microscopy, an image was taken to show the distribution of secretory vesicles (left channel, 488 nm) and the location of peroxisomes (532 nm). The bud as well as other vesicle-dense regions within the mother cell (in the Pex-Boi1 allele) were then bleached to enhance the signal intensity of single secretory vesicles in mother and daughter cells. Single vesicles were tracked utilizing the MOSAIC suite tool plugin for FIJI (see for details in Material and Methods). Single trajectories are color-coded. The edges represent the covered distances, and crosses the positions of each single secretory vesicle over time.
